# Neural and behavioral signatures of the multidimensionality of manipulable object processing

**DOI:** 10.1101/2023.03.29.534804

**Authors:** J. Almeida, A. Fracasso, S. Kristensen, D. Valério, F. Bergström, R. Chakravarthi, Z. Tal, J. Walbrin

## Abstract

Understanding how we recognize everyday objects requires unravelling the variables that govern the way we think about objects and the way in which our representations are organized neurally. A major hypothesis is that the organization of object knowledge follows key object-related dimensions, analogously to how sensory information is organized in the brain. Here, we explored, behaviorally and neurally, the multidimensionality of object processing. We focused on within-domain object information as a proxy for the kinds of object decision tasks we typically engage in our daily lives – e.g., identifying a knife from other types of manipulable objects such as spoons, axes or screwdrivers. To do so, we extracted object-related dimensions from subjective human judgments on a set of objects from a particular object domain – i.e., manipulable objects. We demonstrated that the extracted dimensions are cognitively interpretable – i.e., participants are able to label them; are cognitively relevant for manipulable object processing – i.e., categorization decisions are guided by these dimensions; and are important for the neural organization of knowledge – i.e., they are good predictors of the neural signals elicited by manipulable objects. This shows that multidimensionality is a hallmark of the organization of object knowledge in the brain.

## INTRODUCTION

Our ability to recognize one object amongst many others is one of the most important features of the human mind, and successful completion of our daily activities depends upon it. This is nicely illustrated by the performance of neurological patients that present with semantic impairments, and are severely dysfunctional even when performing mundane activities (1–7). For instance, Patient EW(3) complained of having ‘*no idea about animals at all’* and could not *‘even think about what animals looked like’*, in the context of normal processing of objects from other categories. Our capacity to recognize objects is most probably dependent on how we represent object-knowledge and on how these representations are organized in the brain (8–10). One tenable hypothesis on the organization of object knowledge holds that object representations are, at some level, grounded on, and framed by, relevant object-related dimensions in which objects vary (11–15). These object-related dimensions would allow for the hard, fine-grain, and most times within-category distinctions that are typically required when recognizing objects. However, there are substantial gaps to be closed under such a hypothesis – most importantly, what are these object-related dimensions that govern the finer-grain neural organization of object representations? And how do they guide human behavior? Here, we will extract object-related dimensions from participants’ understanding of objects, and then show that these dimensions are cognitively and neurally relevant.

The mere presentation of an object brings about large amounts of object-specific information (e.g., that a knife has a blade) that come online in an intricate web of object-related knowledge. Moreover, an object can be instantiated by many different exemplars (e.g., my hammer, *Thor’s* hammer), and even a particular exemplar can project an infinite variety of retinal images depending on its location and orientation. So how do we resolve the input from this complex, interconnected, and recursive environment with such ease and proficiency? The answer to this question lies in understanding how object knowledge is represented and organized in the brain such that we can use it efficiently (8–10). Research on the organization of object knowledge in both human and non-human primates points to the existence of clusters of neurons that show categorical preferences for particular object classes such as faces, animals, body parts, places, and manipulable objects (14, 16, 25, 26, 17–24). Moreover, data from brain damaged patients also seems to point to the presence of independent cognitive modules that are dedicated to those same object classes (1–5, 7, 27). Most, if not all, current theories on conceptual knowledge focus on large-scale distinctions such as object category (3, 21, 23, 25, 26, 28), animate/inanimate status (29–31) or other coarse dimensions (32–38) to explain the spatially-specific object preferences within object-specific neural space. For instance, in ventral temporal cortex these object preferences follow a lateral-to-medial organization with clusters of neurons preferring faces or animals situated within lateral fusiform gyrus, and clusters of neurons preferring places or manipulable objects located more medially (10, 39). This organization has been described either as 1) a sharp functional division reflecting true domain distinctions (1, 3, 23, 25, 29, 30, 40–42), laid out, for example, by connectivity constraints between these object-preferring regions and other regions elsewhere in brain that code for the same domain (43–45); or as 2) a continuous map of similarity over one single dimension such as animacy (34), or shape (36, 37) (for other dimensions see (32, 35)).

Thus, most current approaches propose overarching explanations for the general organization of object knowledge by object-preferring regions, in lieu of focusing on the finer-grain organization of conceptual content within these different regions. That is, their focus has been on explaining the pattern of results from neuropsychology and neuroimaging at a general level, appealing to a first principle of organization that has to be somehow, at least indirectly, related with the differences between the domains of stimuli (1, 3, 23, 25, 29, 30, 32, 34–37, 40–45). These efforts, while necessary, may have deterred the field from explaining also the more fine-grain distinctions – those that, in fact, seem to be affected in category-specific brain damaged patients (1–5, 7). Patient EW, as described above, was unable to distinguish *between different members of the domain of animals*, not between stimuli from different domains. Understanding how we identify a hammer from other objects such as axes, flyswatters, or screwdrivers – i.e., those kinds of distinctions that we probably have to deal with every day, and that are affected in category specific-deficits – requires uncovering particular finer-grain types of information than identifying a hammer from a cat, a truck, or a mimosa.

To start, one way to focus on this level of processing is to think of object representations as being organized in particular multidimensional spaces. These spaces preserve individual properties of objects, while reducing the complexity that is inherent to object recognition by situating each object’s representation within key object-related dimensions. This multidimensional space would also potentially retain relative similarity between object representations, as proximity between objects in this representational space would be a proxy of object similarity in the real world (29, 46). Interestingly, organizing information in multidimensional spaces is a rather widespread strategy within the brain. The clearest examples of this organizational principle are found within sensory cortices, with topographical mapping of neural preferences along particular sensory dimensions. For instance, in early visual cortex, multiple dimensions are superimposed on the cortical sheet including polar angle and eccentricity of the location of a stimulus in the visual field/retina, stimulus orientation, stimulus direction of motion, or eye dominance (47–52). The same holds in auditory cortex, where we have several superimposed dimensions that map central properties of sound – e.g., tonotopy, periodicity, loudness, binaural difference (53–58) – or in motor cortex where we may have dimensions related with muscle groups, action types or spatial locations relative to the effectors that coexist within the same motor cortical swath (59, 60).

Recently, there have been some attempts at describing the multidimensional space that underlies object knowledge (15, 61–64). For instance, Hebart and collaborators have suggested a series of dimensions that may explain, at a general level, how individuals represent many different objects. The dimensions obtained are relatively interpretable (e.g., colorful; fire-related) and are able to predict performance in an odd-one-out similarity judgement task. While these efforts certainly advance our understanding on the processes at play during object recognition, they still follow a *“between-category”* overarching approach and sample their stimuli from different categories. As described above, this approach may be at odds with the kinds of complex decisions that humans have to make in their daily lives and that category-specific brain-damaged patients fail at – deciding whether to an object entering their visual field is a hammer or an axe, or whether the animal approaching is cow or a raging bull. Understanding these kinds of fine-grain decisions is probably better served (or at least is also served) by studies that focus on *“within-category”* strategies. Unfortunately, studies probing this level of processing, and that provide behavioral and neural data, are lacking. Here, we implement a principled way of exploring the multidimensional organization of object knowledge, and focus on manipulable objects as a case study in order to restrict our focus on a domain of knowledge and address finer-grain neural and cognitive organization of object knowledge.

We selected manipulable objects as a case study because they have a set of properties that are useful when trying to define object-related dimensions. Namely, 1) they are everyday manmade objects that we perceive and interact with constantly, and are, thus, fairly familiar; 2) they hold relatively defined sets of information associated with them: by definition these objects have particular functions that they fulfill, have associated motor programs for their use, and have specific structural features (e.g., shape) that may help fulfill both their function and facilitate their manipulation; and 3) their visual inspection engages a set of neural regions that includes aspects of the left inferior parietal lobule (IPL), the anterior intraparietal sulcus (aIPS), bilateral superior and posterior parietal cortex (SPL) and caudal IPS, bilateral dorsal occipital regions proximal to V3A, the left posterior middle temporal gyrus and lateral occipito-temporal cortex (pMTG/LOTC), and bilateral medial fusiform gyrus (mFUG; e.g., 16, 20, 70–72, 22–24, 65–69). Moreover, conceptual knowledge about manipulable objects can be impaired (or spared) in the context of spared (or impaired) knowledge about other domains in brain damaged patients (2, 7).

Here, we will demonstrate that a series of object-related dimensions, extracted from participant’s understanding of objects, relates to the neural representations of those objects, and guides object perception. We will first collect subjective similarity measures for 80 manipulable objects. We will independently capture subjective object similarity in terms of their visual properties, the manner with which we manipulate these objects, and the function that is typically associated with them, independently. We selected these knowledge types (i.e., vision, function and manipulation) because these are central for the representation of manipulable objects. We then used multidimensional scaling (73, 74) to identify and extract key dimensions for each of these knowledge types that structure our representational space.

We hypothesize that participants will be able to learn and use these object-related dimensions when recognizing and categorizing manipulable objects. The importance of object-related dimensions to object processing should be analogous to what we know of the importance of sensory dimensions to how we perceive our environment. For instance, perceiving the pitch of a sound is dependent on the tonotopic organization within the auditory system. Tonotopic representations that govern the organization of sound representations in auditory areas are central for extracting the fundamental frequency of a sound, which is itself a necessary step for pitch perception (75, 76). Moreover, retinotopic representations in V1, and their topographical relationships with V5, may be critical for the phenomenon of apparent motion. In the apparent motion phenomenon, two spatially displaced stimuli are presented in alternated fashion to phenomenologically provoke a percept of a stimulus moving between two points. Interestingly, these stimuli elicit retinopically-specific responses in V1 within voxels falling in the illusory motion path (i.e., in areas where no real stimulus was presented 77). These examples demonstrate how the dimensions that organize sensory information in the brain – such as tonotopy or retinotopy – directly impact behavior. Thus, object-related dimensions should also be important when categorizing and identifying objects.

Moreover, we also predict that these dimensions should be able to explain neural responses elicited by manipulable objects in a way that is dependent on the scores of these objects in each dimension. Furthermore, the spatial extent of the neural responses explained by each dimension should strongly relate to the type of content these dimensions represent. Specifically, dimensions extracted from the similarity in the function typically assigned to each object should best explain responses in ventral and lateral temporal cortex as these regions seems to be tuned towards different types of functional uses of objects and actions (78–80); dimensions extracted from the similarity in the manner with which objects are manipulated and interacted with should best explain responses elicited by objects within occipito-parietal and parietal regions as these regions are typically involve in the computation of object grasps and 3D object processing (81–89).This should be especially true for those dimensions whose object interaction and manipulation related content can be directly extractable from the input (such as whether we grasp an object with a precision or a power grasp). Importantly, there is recent data suggesting that those aspects of object interaction and manipulation that are more dependent on knowing what the object is, or from what material it is made of, are more dependent on regions within ventral temporal cortex (66, 90–92) (e.g., grasping an object in accordance with its function). As such, manipulation dimensions that focus more on these aspects may also explain responses within ventral temporal cortex; finally, dimensions extracted from the similarity in how objects look like visually should strongly explain responses in areas within ventral and lateral temporal cortex that seem to be engaged in computing shape and texture properties. In fact, depending on the content of these dimensions, we expect that more lateral temporal areas should be involved in shape processing whereas more medial ventral temporal regions should be involved in dimensions referring to material and texture aspects of the objects (93–95).

Overall, we predict that these dimensions figure critically in how we represent manipulable objects, and potentially support within-category distinctions. Focusing on this level of analysis will thus complement the literature on object recognition that has typically address conceptual knowledge at a between domain level of analysis.

## RESULTS

### Overview

We first selected a set of 80 common manipulable objects (see Table S1 for all the objects; see Methods). These were selected to be representative of the different types of objects that humans use routinely. We then extracted key object-related dimensions that govern our manipulable object representational space. These dimensions represent the building blocks of how we think about and use these objects. One way to unravel such dimensions is to analyze how different objects relate to each other in a large conceptual space (29, 46), and extract the main axes that organize this conceptual multidimensional space. To do so, we obtained similarity spaces for each of the different object-knowledge types tested – visual information, functional information and manipulation information. We presented participants (N=60) with words referring to the 80 manipulable objects and asked them to think about how similar these objects were in each of the knowledge types (e.g., function). We used an object sorting task to derive dissimilarities between our set of objects (Fig. 1A; 74), because sorting tasks have been shown to be a highly efficient and feasible way to obtain object similarities from large sets of objects. We applied non-metric multidimensional scaling (MDS; 73, 96) to these 3 matrices independently to extract key object-related dimensions that structure our object representational space. The selected dimensions were subsequently labelled by a second group of participants (N=43 participants) in order to ascertain their interpretability. Each participant went through all dimensions, and was told that they could generate up to 5 labels per dimension that, in her/his view, best explained the difference between the objects that were situated at the two extremes of the target dimension (see Fig. 1B).

**Figure 1.**
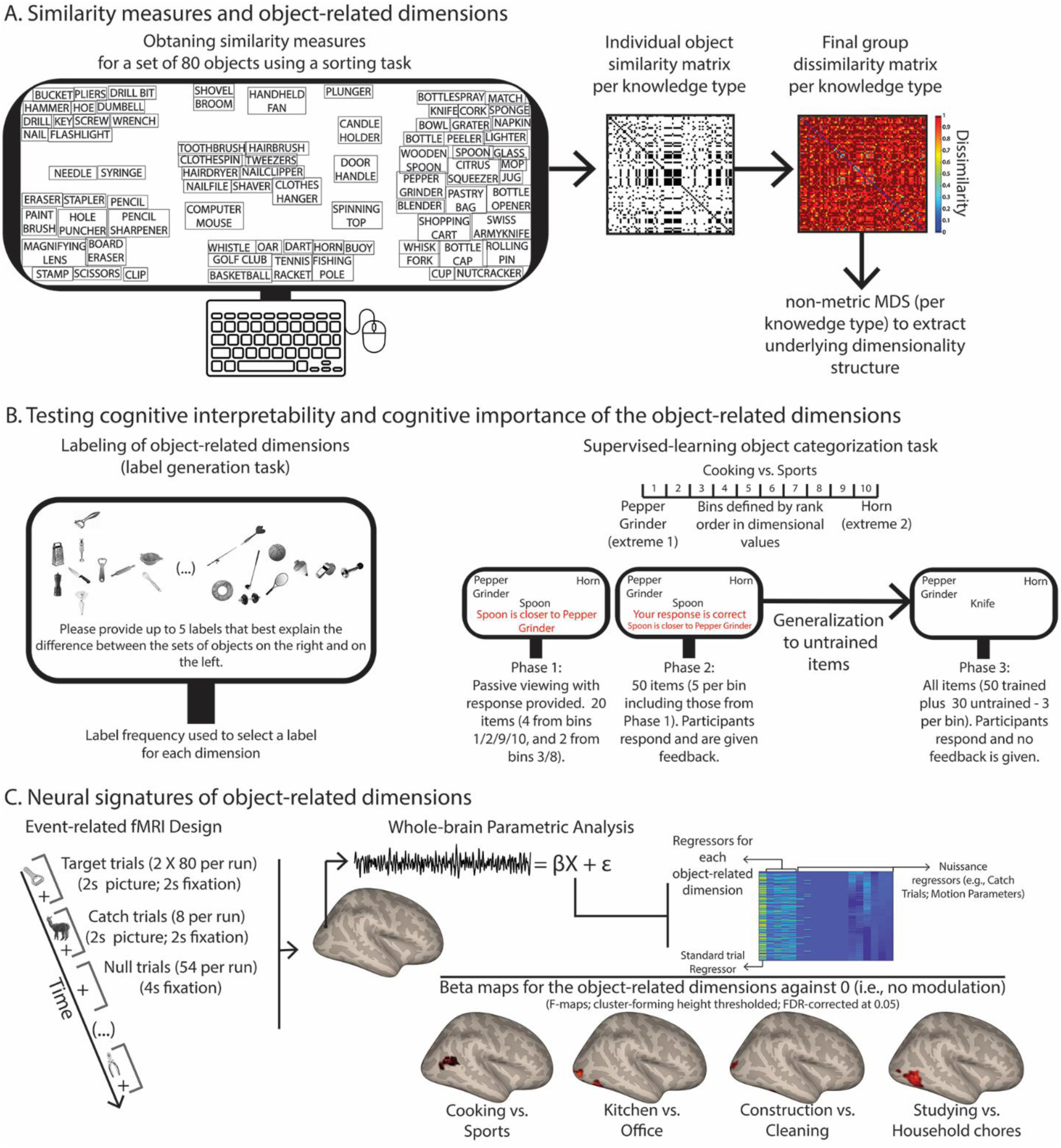
Experimental procedures and analysis pipeline. **A)** In order to extract object-related dimensions we collected similarity measures between our 80 manipulable objects through a pile-sorting experiment. Per individual, we obtained a piling solution for each of the knowledge types (function, manipulation and vision) whereby objects piled together were similar to one another but different from objects in other piles. These piling solutions were coded into dichotomous matrices that represented pile membership. Participant-specific matrices were then averaged and transformed into a dissimilarity matrix (one per knowledge type). Finally, we used non-metric MDS to extract dimensions independently per knowledge type. **B)** We wanted to test whether the obtained object-related dimensions were cognitively important for perceiving and comprehending objects. Firstly, we focused on the interpretability of these dimensions and had a different set of participants perform a label generation task for each dimension. Participants were presented with 20 objects – 10 from each of the extremes of the target dimension – and were asked to provide up to 5 labels that best explained the difference between the two sets of objects. Label frequency was used to select a label for each of the extracted dimensions. We further tested the importance of these object-related dimensions by having yet another set of participants learn to categorize objects according to their scores in each of the dimensions. Participants went through 2 experimental phases where they learned to categorize a subset of the objects in terms of whether they were close to one of the two extremes of a target dimension. Importantly, in a third phase, they were asked to categorize all objects, including a subset of untrained objects. We tested whether participants generalized their learning to untrained items. Percent response performance towards the extreme object with the highest score in the dimension was calculated and fitted with a cumulative Gaussian curve. **C)** Finally, we tested whether the object-related dimensions extracted were able to explain neural responses to objects. We presented the 80 objects in an event-related fMRI experiment using greyscale images, and participants had to categorize each image as either a manipulable object or an animal (the catch trials). We then used parametric mapping to analyze the fMRI data, and tested whether our object-related dimensions could explain the neural responses elicited by the 80 manipulable objects.

Understanding how central these dimensions are in the organization of object knowledge requires, however, establishing that they affect and predict behavior and neural responses to objects. To this end, we tested whether these dimensions could guide participant’s behaviors towards target manipulable objects. We recruited a new group of participants (N= 210; 10 per dimension, and 60 for control dimensions) to perform a supervised-learning categorization task. In this task, we taught participants to categorize a subset of our 80 objects in terms of their scores along a target dimension, and we tested whether their learning could be generalized to a subset of untrained items. That is, can our object-related dimensions be implicitly learned and guide the behavior of a set of naïve participants? In Figure 1B, we show the different phases of this experiment – in the first two phases of the experiment, participants were taught to categorize a subset of objects as to whether they were closer to the object in one or the other extreme of the target dimension and were given clear feedback as to which response was correct. In the third phase of the experiment participants were required to categorize trained and, importantly, untrained items and were not given any feedback on their response (see Methods). All key object-related dimensions were tested in this experiment (each participant was tested on only one dimension).

Moreover, we added two control dimensions. In one of these controls, we took one of the real dimensions and randomly shuffled the scores of the dimension for the individual objects. For a more stringent control, we took lexical frequency values (97) – i.e., count of the times a particular lexical entry appears in a text corpus per million – for each of the objects and rank ordered them in terms of these values. We used these dimensions to control for reliable generalization of object-related dimensional learning to untrained items. We used a training task so as to implicitly prime participants to the target dimensions – as there were several dimensions per knowledge type that could be used by the participants if we opted for an untrained experimental situation. Note that, implementing the kinds of controls described above demonstrates the learning is not unspecific and is focused on the dimensions extracted.

Finally, we needed to show that the extracted dimensions were able to explain neural responses to manipulable objects – that is, to demonstrate that the organization of our neural representations about objects also relies on this multidimensional space. To test this, we developed an event-related functional magnetic resonance imaging experiment (fMRI; see Fig. 1C) where we presented a new group of participants (N=26) with images of the 80 manipulable objects (see Figure S1 for examples of the images used in this experiment). We asked whether the scores of each object in each dimension were able to explain neural responses elicited by those same objects. We used parametric analysis (98–100) over the fMRI data, and cast our key dimensions as first-order (i.e., linear) parametric modulators in a General Linear Model (GLM; see Fig. 1C). That is, for each stimulus in the design matrix, the corresponding dimensional scores were assigned as modulation values. For each participant we obtained a set of beta maps for each dimension, where voxel intensities describe the extent to which stimulus responses vary as a function of their score along a given dimension. Group-level F-test maps (cluster-forming height-thresholding at p<.001 – see exceptions below; FDR corrected at *p*<.05) for each key dimension against zero (i.e., against no modulation) were computed.

### Extraction and labelling of key object-related dimensions

The number of dimensions in the final MDS solution per knowledge type was determined based on *stress* value (Kruskal’s normalized stress; 96). The final solutions led to the extraction of 4 orthogonal functional dimensions (stress value for the solution 0.09), 6 orthogonal manipulation dimensions (stress value for the solution 0.08), and 5 orthogonal visual dimensions (stress value for the solution 0.09; see Fig. 2 for these dimensions; see Figure S2 for scree plots; see Methods for details).

**Figure 2.**
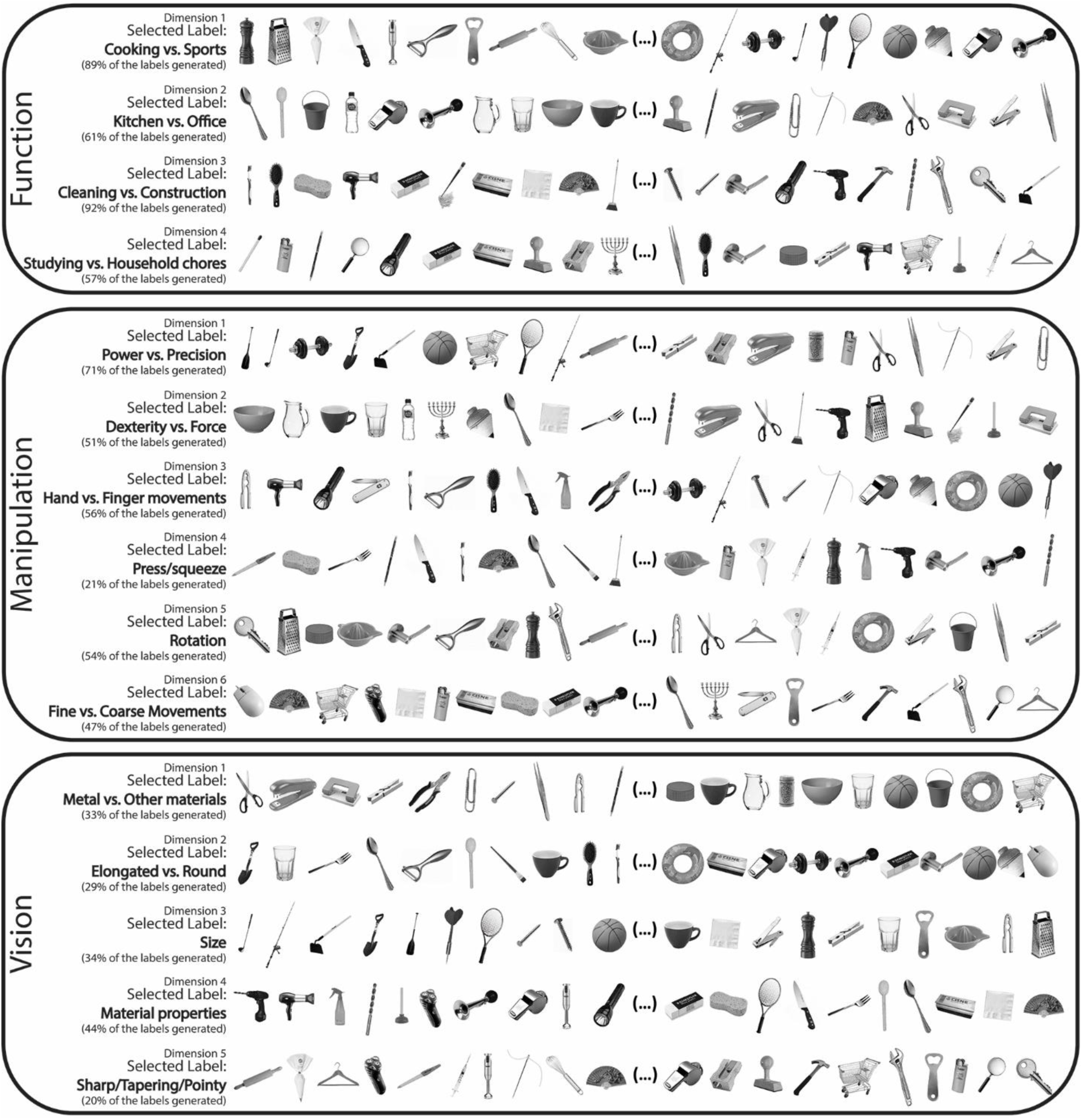
Interpretability of the objected-related dimensions. Here, we present the selected 15 dimensions that govern the internal representation of our set of manipulable objects. Per dimension, we present the 20 objects with the most extreme scores (10 from each extreme). Labels were then selected based on the frequency of label generation by the participants. On the left side of each dimension, we present the collated selected label, as well as the percent generation score for that label (see Figure S3 for the label frequency plots).

Thus, by reducing the dimensionality of our similarity measures, we were able to extract a limited set of key object-related dimensions that govern our understanding of a large set of objects. However, an initial test of our dimensionality solution is to understand how interpretable these dimensions are in terms of their relationship with the objects. We collected and collated labels for each of the extracted dimensions. In Figure 2, we present a rank order distribution of the 20 most extreme objects per dimension (10 from each extreme), along with the selected label and the collated percent label generation score (a visualization of the labels generated can be found in Figure S3). For most dimensions, participants reliably reached similar interpretations. The most consensual labels generated for the visual dimensions include aspects related with shape (e.g., roundedness), size, or material properties (e.g., presence of metal); the labels for the functional dimensions include aspects related to different types of activities performed (e.g., cleaning vs construction), or the context in which the objects are seen (e.g., kitchen vs. office); and labels for the manipulation dimensions include aspects related to grasp types (e.g., power vs. precision), different types of motion (e.g., rotation), and object properties that relate downstream to object manipulation (e.g., need for force or dexterity). There are some dimensions whose labels were not consensual (e.g., selected labels for dimensions 4 of manipulation and 5 of vision present label generation percent scores of about 20%). Interestingly, these lower scores are observed more consistently in later (e.g., the fifth dimension) rather than earlier (e.g., the first dimension) dimensions, as this may relate to the fact that these dimensions explain less of the total variance. Overall, however, the reliability of the labels attributed to each dimension seems, as a first pass analysis, a demonstration of the feasibility of our approach, and the centrality of these dimensions to the processing of objects.

### Object-related dimensions guide object categorization behavior

A much stronger test of the centrality of these dimensions is to understand whether and how they can guide behavior. Figure 3 shows categorization performance on untrained manipulable objects, as to whether the target objects were more categorizable as closer to one or the other extreme object in a target dimension. Cruciallly, participants were able to generalize their learning on trained objects to untrained objects – categorization of untrained manipulable objects was affected by their dimensional score on the trained dimension. This is true for most key dimensions, but not for the control conditions (random distribution and lexical frequency).

**Figure 3.**
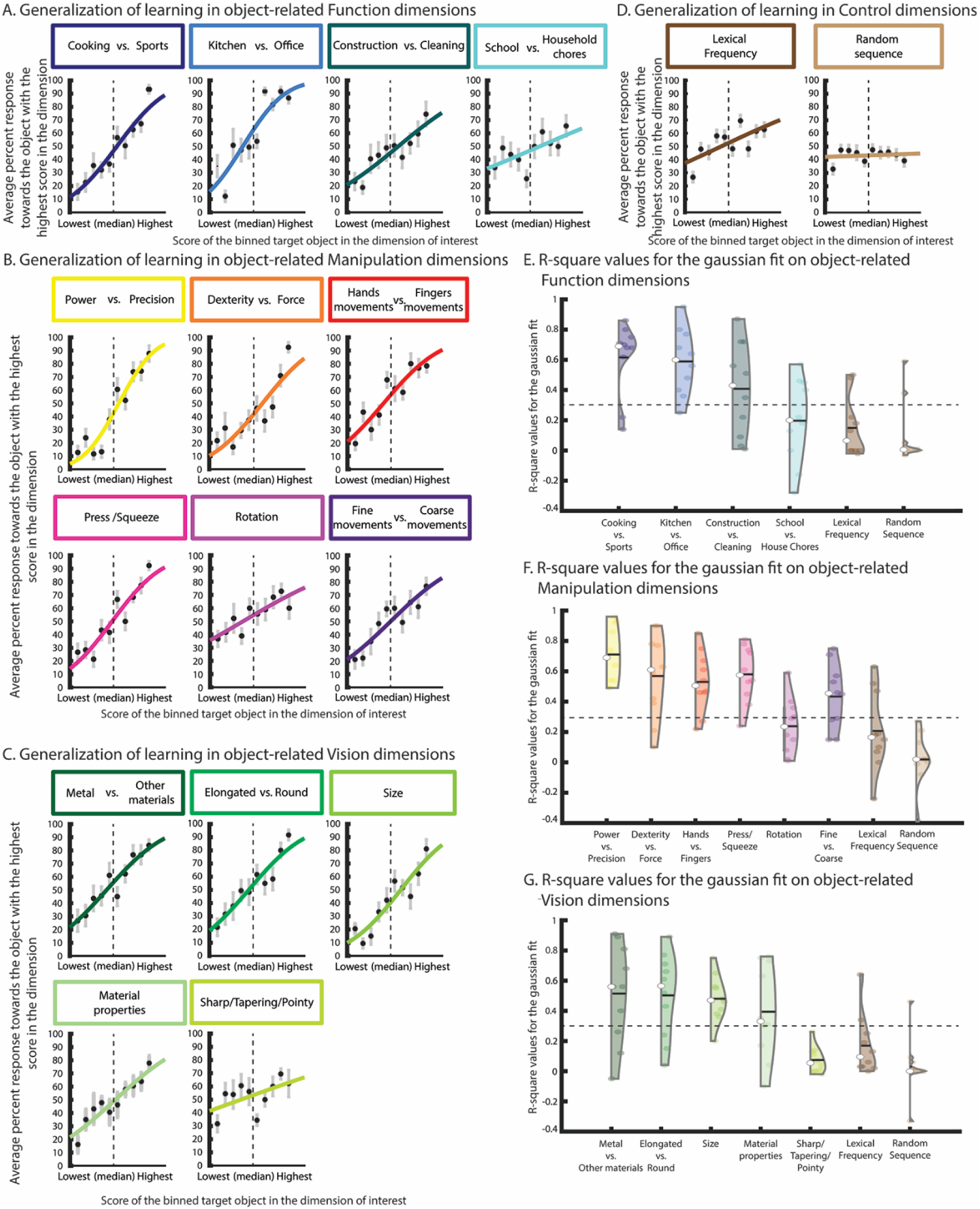
Supervised learning of object-related dimensions in a categorization task. A-D) Here we show percent response towards the object with the highest score for each of the object-related dimensions and the two control dimensions (i.e., towards the extreme with the highest score). Percent responses were averaged within each of the ten bins, and a cumulative Gaussian curve was fitted on the data of each individual. The presented plots are based on the average of all participants. Error bars correspond to SEM; depicted cumulative Gaussian curve was fit on the average percent results for visualization purposes; **E-G)** Violin plots of the R-square values of the Gaussian fit for each dimension for each participant. Dashed line corresponds to R-square = 0.3.

To analyze these data, we first averaged responses per bin (see Methods for the definition of the bins). We then fitted a cumulative Gaussian curve to the bin-specific percent categorization responses towards the object with the highest score in the target dimension. We expected that if the object-dimensions were cognitively important for our ability to process and recognize objects, then participants would be able to generalize their learning of the target dimensions to untrained items – thus, percent responses towards the extreme object with the highest score should increase as a function of the increase in the dimensional score per bin. R-square values per participant were obtained as a measure of goodness of fit between the cumulative Gaussian curve and each participant’s data.

As can be seen in the violin plots in Figure 3, for the key object-related dimensions, most participants presented R-square values well above 0.30. This specific cutoff is necessarily arbitrary, but we used this liberal criterion to allow us to determine if any given dimension is reasonably predictive of each participants’ behavior. That is, whether it contributes to the representation of the object as well as guide behavior. It is particularly impressive that most of the extracted dimensions have high explanatory powers, indexed by high R-square values. However, there was a trend, for lower goodness of fit for those dimensions there were lower in the stress ranking of our MDS solution (the last dimensions of function and vision and the second to last dimension of manipulation). This reduced generalization of learning for dimensions with lower values of stress is actually a demonstration of the specificity of this learning to object-related dimensions and not to any dimension. Even with such a liberal criterion it was clear that the control conditions did not explain the behavioral data, with R-square values consistently below 0.3. As such, this is a strong test of the centrality of these dimensions in organizing our representational space, and of whether these dimensions may generalize beyond the individuals that generated them.

### Object related-dimensions predict neural responses elicited by manipulable objects

Our categorization experiment showed that the MDS-extracted object-related dimensions can guide object processing and be central to our object recognition efforts. A final stage in demonstrating that multidimensionality is a signature of high-level object processing, as it is of low-level sensory processing, is to demonstrate that the organization of our neural representations about objects also relies on this multidimensional space.

As can be seen in Figure 4, our object-related dimensions can account for neural signal elicited by viewing manipulable objects. That is, all key object-related dimensions show linear slope (beta) values that significantly differ from zero – i.e., all dimensions significantly modulate the neural responses to manipulable objects in different regions as a function of the object dimensional values. Note that the function dimension “Cooking Vs. Sports” shows limited coverage under a cluster-forming height-thresholding of p<.001, and the dimension “Material properties” fails to survive correction at that level. Nevertheless, these two dimensions show significant results under a slightly more lenient cluster-forming height-thresholding of p<.005 (for individual maps of all dimensions see Figure S4).

**Figure 4.**
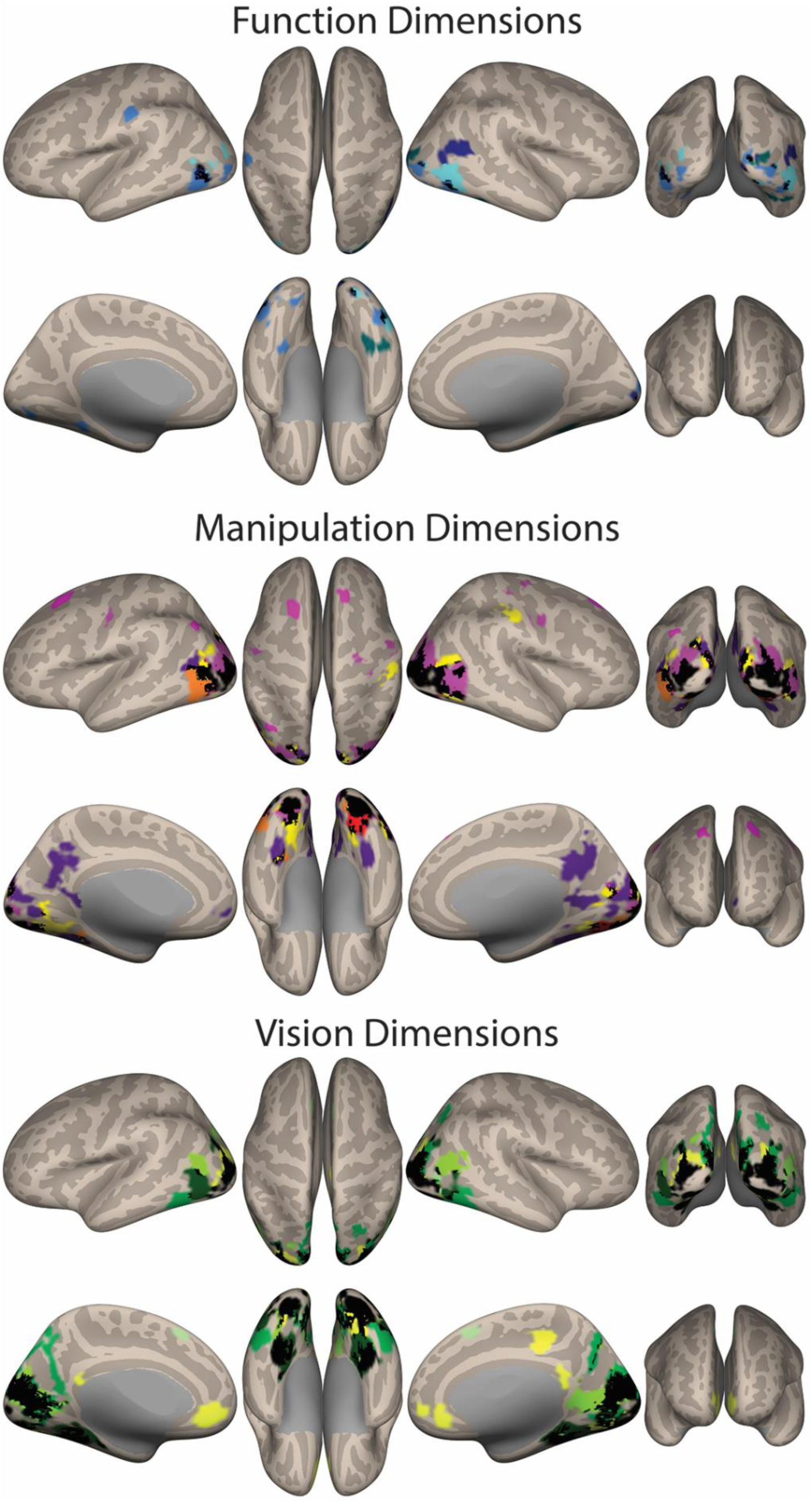
Neural effects of the object-related dimensions. Here, we show an overlap map with all object-related dimension (F-maps per dimension against zero – i.e., against no modulation) per knowledge type, each dimension color coded by the colors in Figure 3. Black corresponds to areas where at least two individual dimensions overlap (all individual F-maps cluster-forming height-thresholded at p<.001 – except for the dimensions “Cooking vs. Sports” and “Material properties” that are cluster-forming height-thresholded at p<.005 – and all corrected at FDR p<.05).

Moreover, these object-related dimensions appear to capture signal variability in particular regions, mostly within those regions that show a preference for manipulable objects (16, 20, 69– 72, 22–24, 45, 65–68) – that is, many of these dimensions overlap spatially in particular regions. This is in line with the role of dimensionality in explaining neural organizing of information: perhaps in the same way as the different dimensions that rule the organization of low-level sensory-motor cortices overlap spatially, so do dimensions that rule the organization of object knowledge in the brain.

These data then suggest that multidimensionality is a signature of cortical processing and neural informational representation and organization at multiple levels of abstraction and complexity.

### Object related-dimensions generally code for knowledge-type (vision, manipulation, function) specific information in the brain

Interestingly, and as predicted, these results show specificity as a function of the type of content these dimensions represent. This can be seen more generally when we observe the neural parametric maps of these dimensions grouped by knowledge type. For instance, as a group, function dimensions seem to collectively explain responses (i.e., their responses overlap or are in proximity of each other) in pMTG and lateral occipital cortex (LOC; see Fig. 4). Remarkably, these regions have consistently been shown to be associated with object-related action knowledge and meaning (78, 79, 101, 102), and to respond topographically to different types of action (80) and to how objects interact to fulfill the intentions of an actor (103). As such, they are potentially central for the processing of object-related function information.

Manipulation dimensions show a lot more overlap within occipito-parietal regions (in the vicinity of V3A, caudal IPS, and Precuneus), most probably due to the fact that these dorsal occipital regions are involved in object-specific 3D processing (27, 81, 111, 82, 104–110), and participate in object grasping (83–89). These manipulation dimensions also overlap in medial ventral temporal cortical regions, posterior lingual gyrus and pMTG/LOTC. pMTG/LOTC, as described above, has been consistently associated with object-related action (78, 80), and shows sensitivity to different types of actions and how appropriate an object grasp is (66, 90). Moreover, the involvement of ventral temporal areas in action has been recently demonstrated for object-related action observation and execution. This involvement seems to be related to computing object properties that impact action but that have to be derived from visual and functional processing of objects (e.g., and object’s weight, how objects are grasped to perform their function 43, 68, 91, 92, 112–114).

Finally, vision dimensions show major overlap in ventral temporal cortex and posterior lingual gyrus, as well as in lateral temporal cortex, in line with the involvement of these regions in the processing of form and of surface texture and material properties (93–95). These dimensions also overlap in posterior dorsal occipital regions, potentially as consequence of high-level object-related processing that seems to be also happening within these regions (111, 115), as well as the computation of 3D representations (81, 82, 106–110, 116, 117).

### Object related-dimensions maintain content specificity in their neural responses

Specificity in the multidimensional organization of object knowledge can also be observed when we look at these dimensions individually (individual parametric F-maps against 0 – i.e., no modulation – can be seen in Figure S4; The F-maps for the first two dimensions of each knowledge type are presented in Figure 5). These maps show particular aspects that relate to the content represented by each of these dimensions.

**Figure 5.**
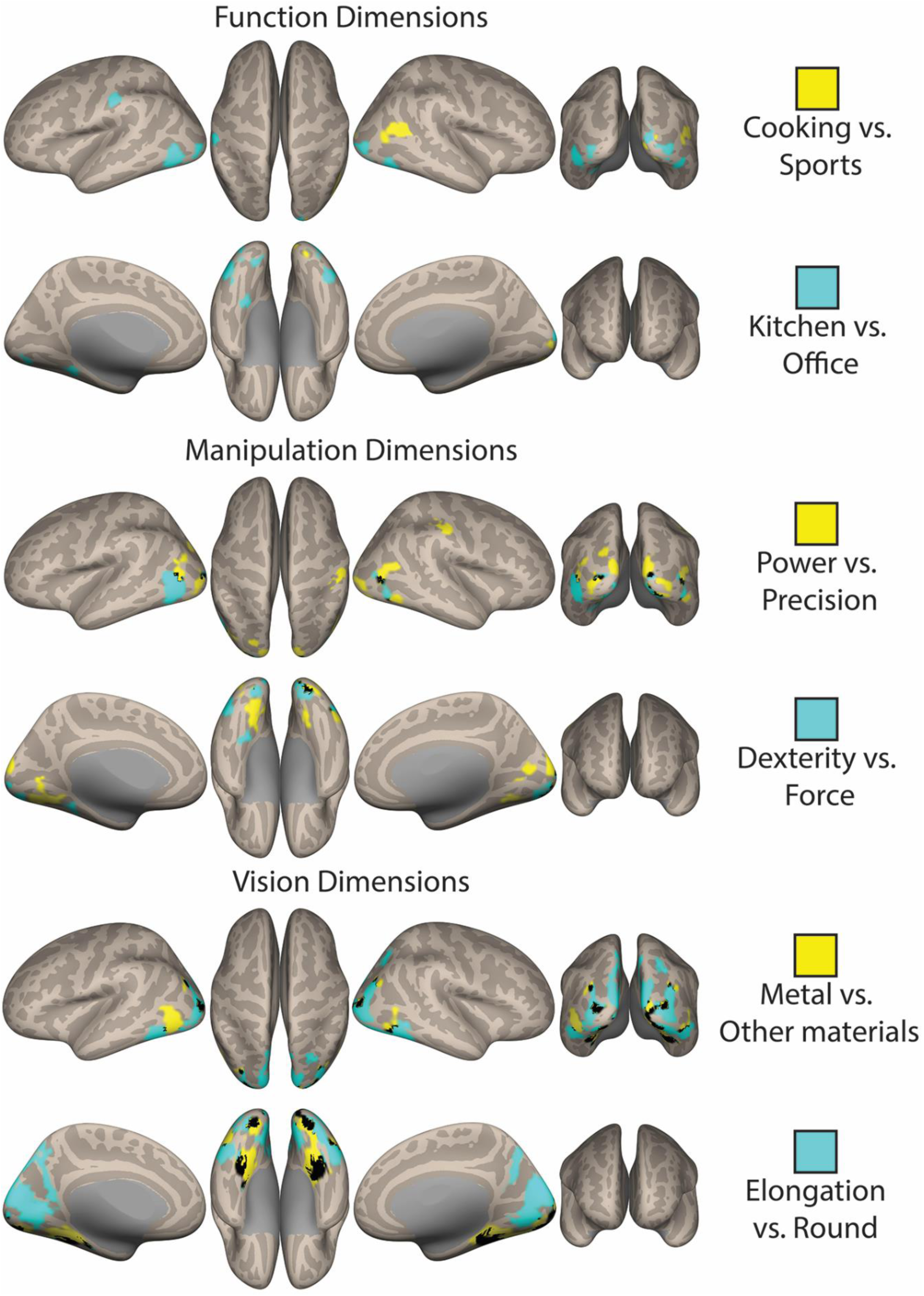
Content specificity in the neural responses. Here, we show modulatory effects (i.e., the beta values of each dimension that are significantly different from zero) of the first two dimensions of each knowledge type (all individual F-maps cluster-forming height-thresholded at p<.001 – except for the dimension “Cooking vs. Sports that is height-thresholded at p<.005 – and corrected at FDR p<.05). In yellow we present the first dimensions of each knowledge type (“Cooking vs. Sports”; “Power vs. Precision”; “Metal vs. Other Materials”), whereas in light blue we present the second dimensions of each knowledge type (“Kitchen vs. Office”; “Dexterity vs. Force”; “Elongation vs. Round”). Each maps shows voxels whose signal is explained by the dimension when compared to no modulation (i.e., when compared to 0). Black corresponds to areas of overlap between the two dimensions presented per map.

The first two function dimensions (“Cooking vs. Sports” and “Kitchen vs. Office”) are able to explain neural responses within broader pMTG/LOTC, although the dimension that relates to the physical context in which the objects are typically observed (“Kitchen vs. Office”) explained activations in more inferior aspects, whereas the dimension that relates to particular types of activity (“Cooking vs. Sports”) explained more superior activation (see Figure 5). Interestingly, pMTG/LOTC has been shown to be topographically organized in a superior-to-inferior gradient that codes for sociability versus transitivity (80) – cooking and playing sports may reflect more action-related sociability than the context in which actions are performed (Kitchen or an office). Moreover, the dimension “Cooking vs. Sports” significantly explains activation in a small area within right lingual gyrus extending superiorly to the cuneus. Interestingly, studies that focus on the mere presence of other objects during transitive action (i.e., contextual objects), and the amount with which these objects relate to the goal of the actor, lead to activations within the cuneus (among other areas) around the regions obtained here (103, 118, 119). Finally, the dimension “Kitchen vs. Office” explains signal also within the border between left IPL and post-central gyrus, and, importantly, within the parahippocampal gyrus, centered on the coordinates typically shown for the parahippocampal place area (PPA; 26, 41). PPA has been repeatedly shown to be related with processing of spatial relations, layout, navigation, and different (visual) environments – potentially what differentiates a kitchen and an office.

The two selected manipulation dimensions represent different content. While both relate to aspects that are important for manipulating an object, one of them relates to properties of objects that are directly retrievable from the visual inspection of the object (“Power vs. Precision”; i.e., what kinds of grasps are afforded by the shape of a manipulable object), the other relates to properties that are obtained indirectly from the inspection of an object – perhaps from an object’s surface texture and/or material properties (“Force vs. Dexterity”) – and that potentially relates to the computation of weight. This difference seems to be mirrored in the parametric maps obtained for each dimension (Figure 5). Overall, the “Power vs. Precision” dimension is capable of explaining activity within regions of the posterior parietal cortex – namely aIPS (extending to post-central gyrus), at or in the proximity of regions shown to be associated with hand shaping for grasp, and different types of grasps (43, 71, 88, 120), and regions proximal to V3A that seem to code for shape properties and 3D representations of objects independently of action (81, 95, 106– 108, 111, 121, 122), as well as grasp planning and different grip formations (i.e., power, precision; e.g., 33, 43, 125, 83, 87–89, 114, 120, 123, 124, but see e.g., 83, 84; see Figure 5). The dimension “Force vs Dexterity”, shows no occipito-parietal and posterior parietal activations, but shows bilateral ventral (both anterior within lingual gyrus, and posterior within parahippocampal gyrus) and lateral activation (see Figure 5), consistent with the involvement of surface and material properties in deriving the amount of force or dexterity needed to act with these objects. Importantly, the representation of grip force(112), and object weight – an object property that allows for the derivation of the amount of force to apply to an object(91) – engage posterior fusiform, LOC and areas linked with texture processing within the ventral temporal cortex (in the proximity of the regions obtained here).

The two selected visual dimensions also relate to a major distinction in visual processing – that of dimensions that encode content related with material and surface properties (e.g., “Metal vs other materials”; see also “Material properties”) versus those that encode content related with shape and visual form (e.g., “Elongated vs. Round”; see also “Size”). The dimension “Metal vs. other materials” is related with material and surface properties of the objects. Interestingly, the largest clusters of activation for the dimension “Metal vs. other materials” are medially along the collateral sulcus (posterior to anterior) bilaterally, parahippocampal gyrus and the posterior lingual gyrus (see Figure 5). These regions have been associated with the processing of material and surface properties of objects (93–95, 126). The dimension “Elongated vs. Round” shows a major cluster of explained signal in lateral ventral cortex – a set of regions that are a center for shape processing, and whose disruption severely affects shape processing (93–95, 117, 127, 128) – as well as in inferior occipital cortex (see Figure 5), and also strongly in some aspects of dorsal occipital cortex (see Figure 5). These regions are exactly those that have been shown to be coding for object roundedness and elongation (65, 104, 114, 122, 129).

Overall, then, our results show that neural responses elicited by manipulable objects are explainable by their scores on key object-related dimensions particularly within regions that typically prefer manipulable objects to other categories of objects (e.g., faces). Moreover, the ability of these dimensions to explain neural responses to objects seems to be related with the kinds content that the dimensions represent.

## DISCUSSION

Unravelling the organization of object knowledge in the brain is a necessary step in understanding one of the most ubiquitous processes implemented by the human mind – that of recognizing objects – making it a central endeavor for the cognitive sciences(8–10). Importantly, most efforts to understand object recognition and the organization and representation of conceptual information in the brain, have focused on explaining between category differences, or on finding an overarching explanation for object representation. However, the available neuroimaging and neuropsychological data (1–5, 7, 14, 17–27) seem to point, at least, to the parallel need to also look into more fine-grain distinctions – i.e., into within-category organization of information. Here, we tested a principled way to explore the multidimensionality of object processing focusing on the category of manipulable objects. Specifically, we extracted object-related dimensions from human subjective judgments on a large set of manipulable objects. We did so over several knowledge types (i.e., vision, function and manipulation), whose selection was motivated by the typical characteristics that best describe these objects. Our results demonstrate that these dimensions are cognitively interpretable, that they guide our ability to categorize and think about objects, and that they explain neural responses elicited by the mere visual presentation of these objects. Moreover, our results show that these dimensions are highly generalizable across individuals – all participants were naïve, were independently selected for each experiment, and were not aware, *a priori*, of the objects that they would be presented with nor which dimensions were being tested – and even across modalities of presentation of the stimuli (words vs. pictures), suggesting some level of universality.

Our data shows that the way in which we perceive and think about manipulable objects can be captured and governed by the object-related dimensions extracted from our subjective understanding of the world. Importantly, we trained participants to categorize objects while implicitly using a metric based on (one of) these dimensions. Critically, we showed that their learning could be generalized to untrained items – that is, participants were able to categorize items that they had never seen based on their understanding of how these novel objects scaled with the dimension they were implicitly taught. The generalization results were dramatically different when the dimension to be learned was non-object related (either lexical frequency or a random shuffle of real dimensional values) – in these cases no generalization to untrained items was observed. Note that these participants were completely naïve to all the major experimental manipulations – they did not know what the dimensions to be tested were; they did not know that there were 80 manipulable objects in total; and they did not participate in any of the other experiments. These data strongly suggest the validity of these dimensions in organizing the conceptual space of manipulable objects, as well as of our approach to understanding the multidimensionality of object space.

Moreover, we also show that these same dimensions can explain neural responses elicited by our set of manipulable objects – that is, these dimensions show explanatory power not only at the behavioral level but also at the neural level. Several results are worth discussing here. Firstly, variation in the scores of each object in the target dimensions explains neural signals elicited by the mere presence of those objects. This is a demonstration of the importance of the extracted object-related dimensions in the process of representing and recognizing objects. Secondly, our object-related dimensions seem to explain, collectively, neural responses mainly within areas that typically show preferences for manipulable objects when compared to other categories of objects (16, 20, 70–72, 22–24, 45, 65, 66, 68, 69). This overlap likely mirrors the fact that we investigated the finer-grain object-specific representations within manipulable object preferring regions. That is, we are going beyond describing large principles of organization (e.g., domain, animacy; e.g.,3, 21, 34–38, 23, 25, 26, 28–32) that explain the macro-organization of large swathes of cortex, and focus on the finer-grain organization of representations within the divisions proposed at the macroscopic-level(1). Thirdly, beyond the general overlap described above, there is also specificity in how these dimensions, individually, explain neural responses to manipulable objects. That is, each dimension captures specific aspects of the processing of manipulable objects, suggesting a distributed approach, within the macroscopic within-category organization(1), that codes for object properties constituent of an object’s conceptual representation. It is also noteworthy that content differences between dimensions led to clear neural dissociations. For instance, the first two visual dimensions (and the remaining ones) refer to a major divide in visual processing – that of processing of texture and material properties versus processing of shape and form properties. This major divide was mirrored in how these two dimensions explained neural responses in different parts of the cortical surface following our initial expectation – the dimension that referred more to material properties (“metal vs. other materials”) explained neural response to objects within more medial ventral temporal cortex, whereas the dimension that referred more to visual form (“elongated vs. round”) explained more lateral aspects of the fusiform gyrus and lateral occipital areas. Fourthly, and interestingly, the specificity of these dimensions may actually relate to the kinds of mid-level object properties (e.g., grasp-type within dorsal occipital cortex; metal versus other materials within ventral temporal cortex; object elongation within LOC and occipito-parietal aspects) that have been proposed to constitute a crucial stage during object recognition (130–133).

Furthermore, there seem to be major processing centers that display truly multidimensional representational spaces for manipulable objects. Specifically, neural signal in pMTG/LOTC, the collateral sulcus and parahippocampal gyrus, dorsal occipital cortex in vicinity of V3A, and IPS is explained by several of the object-related dimensions that we extracted. This suggests that analogously to the organization of sensory-motor cortices by multiple (at least relatively) independent dimensions in tandem (e.g., the organization of visual information by polar angle, eccentricity, stimulus orientation, and stimulus direction of motion in early visual cortex; e.g.,47– 50, 52), multidimensionality is also foundational to organizing object information and object representations in the brain. The implication, then, is that multidimensionality is a major property of neural and cognitive processing, and a central principle of organization of our object-knowledge space: it is present chiefly in sensory-motor representations and processes (47–49, 51, 52), but also, and importantly, in the processing of complex objects.

Importantly, not only is our data in line with what has been obtained when trying to understand overall multidimensionality of object processing (15, 61, 134), but they also further our understanding of the fine-grain properties of object processing that was lacking hitherto. Several approaches have demonstrated, as we have here, that multidimensionality is central for object processing. For instance, Hebart and collaborators (15), Fernandino and collaborators (64) and Huth and collaborators (61) have all suggested a series of dimensions, experiential features or general principal components that underlie object representations. Interestingly, when we look at some of the individual dimensions proposed by these groups (e.g., round; sport-related; 15) they do relate to those that we of have obtained here. Moreover, the nature of the dimensions reported here, as described above, potentially relates to mid-level object properties – and thus are probably similar to what Fernandino and collaborators (64) call “experiential features” (notwithstanding potential differences in kind of format that we and Fernandino and collaborators advocate for these representations; e.g., modal or amodal). Furthermore, the ability of our dimensions to explain neural responses is most appreciable within associative cortex, in line with what was shown by these researchers (64). However, one aspect that differentiates these previous research efforts and what we present here is, in fact, that all of these previous efforts have focused on a between-category approach, in lieu of the within-category approach we use here. Interestingly, some (perhaps even most, in some cases) of the dimensions obtained by these authors do relate to domain (e.g., Tool-related; Animal-related; Body-part-related; (15)). This brings about two central issues: 1) that, in fact, domain seems to be a major principle of organization; and 2) that the presence of these categorical dimensions – dimensions that probably do not conform to a continuous but rather to a discrete organization – may exhaust much of the variance present in the data (e.g.,15, 61, 64). Overall, these issues call for a within-domain scrutiny of the multidimensionality of mental representations and object processing. Here, we went further and explored the more fine-grain details of object processing at a within-category level, and showed content-specific dimensions that guide the way we perceive manipulable objects beyond (and in a way independently of) the macroscopic differences between domains. These content-specific dimensions are central to the representations we build of the objects we perceive. We can then use these representations in the process of identifying objects and compare them to the object representations we have stored in our long-term memory.

The spatial extent of the cortical regions whose responses are explained by our dimensions merits some discussion regards. Specifically, the dimensions obtained here explain responses primarily within cortical regions that could be considered relatively posterior, and potentially visual (but certainly associative cortex). Note, however, that for instance functional dimensions explain responses in pMTG and ventral temporal cortex. Although these regions could be still be considered visual, the nature of their representations are hardly vision-specific and certainly not low-level (78–80, 101–103). For instance, these same regions are engaged by the auditory presentation of manipulable objects in the congenitally blind, undoubtedly showing that these representations cannot be purely visual (39). Moreover, all of the manipulation dimensions that explain, mainly, ventral temporal cortical regions refer to aspects of manipulation that are not clearly associated with actual visuomotor processing, but rather with processing that will then have an effect on how we interact and manipulate an object (e.g., calculating weight for determining amount of force). Interestingly, it has been shown that regions within ventral and lateral temporal cortex are engaged in the processing of these kinds of information (66, 91, 92). The remaining manipulation dimensions strongly explain dorsal occipital regions, but also more anterior parietal areas and pre-frontal regions, as expected given the involvement of these regions on visuomotor computations (e.g., first manipulation dimension explains responses in aIPS). Nevertheless, our dimensions do explain responses in relatively posterior areas. It is important to mention that these more posterior regions do support object processing of particular knowledge contents that are not low-level and strictly visual – e.g., material and surface properties in lingual gyrus 93–95; grasping in V3A 83–89, etc. Notwithstanding, one possible explanation for these more posterior results may be that our task and trial structure are more inducive of responses in putative visual cortex – objects are presented visually and we include changes from object to fixation that may ramp up contrast responses, which may lead to stronger results in visual areas. Note, however, that we have included in our GLM a predictor that models all the stimuli, which should account for low-level contrast-based responses, suggesting that our results are not influenced by such low-level confounds.

One outstanding question in terms of the generalization of this approach, is whether these dimensions are uniquely capable of explaining neural signatures of, and guide behavior towards, manipulable objects, or whether they are sufficiently explanatory of information from other domains of stimuli. One possible intermediate position may be that these dimensions might be explicative of content that engages the neural patches that are chiefly engaged by manipulable objects. Thus, these dimensions could then partially explain neural signal elicited by other kinds of stimuli, perhaps to the extent that those stimuli also minimally engage those patches. For instance, it is possible that the manipulation dimensions may also explain certain neural and behavioral patterns associated with the domain of fruits and vegetables – a melon is heavier than a raspberry, and different levels of force and dexterity will be required when manipulating these fruits; a grape and a cucumber will require different kinds of grasps when we interact with them.

Another potential outstanding question relates to how these dimensions fit with cultural differences, and the phylogenetic environments the human species went through. Perhaps some of the labels obtained for the object-related dimensions are very contemporary, fitting a modern way of living (e.g., Kitchen vs. Office). Others, however, might be important irrespective of the temporal and spatial location of an individual (e.g., Elongation, Power vs. Precision). There are potentially two ways in which multidimensionality of the organization of object knowledge might be immune to the cultural and phylogenetic environment. One would lean on the plastic nature of our brain in order to use different dimensions in a manner highly dependent of learning possibilities. This might be analogous to modern literate humans repurposing parts of left Fusiform gyrus to represent words, which region would likely have been used to represent faces in our illiterate ancestors (135). Another possibility would suggest that these dimensions are (temporally and spatially) universal, but the labels that humans generate for these dimensions might be relatively crude and would plausibly be different through times and cultures. Speculatively, the latter may be favored. The manipulation and visual dimensions uncovered here seem to code aspects of objects that are central to the processing of visual stimuli and our interactions with “things” in general – be it man-made objects, wood sticks, or flint stones. These dimensions would thus be suitable to the processing of stimuli across cultures and/or phylogenetic times. The functional dimensions that were uncovered may, at first sight, be harder to reconcile with an evolutionary and cultural perspective. However, core aspects of sociability (Cooking, Sports), working, and hunting and gathering (Office; house chores), and constructing are probably essential functional goals that might be relatively stable across cultures and phylogenetic environments. Thus, it is highly possible that the dimensions uncovered, although crudely labelled by our participants, may be fundamental dimensions in the organization of manipulable objects.

Overall, then, we show that object-related dimensions, extracted from the subjective understanding of a large group of individuals over a large set of manipulable objects, can guide our behavior towards objects and can explain object-specific neural responses, as well as finer-grain organization of object-content in the brain, suggesting that multidimensionality is a hallmark of neural and conceptual organization.

### Methods

#### Participants

A total of 339 individuals (305 women) from the community of the University of Coimbra participated in the experiments (age range: 18-41): 60 in the object similarity sorting task, 43 in the label generation task (23 were presented with words and 20 with pictures), 270 in the supervised learning categorization task, and 26 in the fMRI task. The size of each sample corresponds to typical samples sizes of these kinds of experiments (e.g., 44). All experiments were approved by the ethics committee of the Faculty of Psychology and Educational Sciences of the University of Coimbra, and followed all ethical guidelines. Moreover, participants provided written informed consent, and were compensated for their time by receiving either course credit on a major psychology course or financial compensation.

#### Stimuli

We first selected a set of 80 common manipulable objects (see Table S1 for all the objects; see Figure S1 for examples of the images used). These were selected to be representative of the different types of manipulable objects that we use routinely. For Experiments 1 and 3 we used words to represent the object concepts, whereas for Experiment 4 we used images. We used both images and words separately for Experiment 2. Images were selected from the world wide web. We used an imaging editing software to crop the images, extract any background, and gray scale and resize the images to a 400-by-400 pixels square. Presented images subtended approximately 10° of the visual angle. We selected 10 exemplars per object type in a total of 800 images. For Experiment 4, we additionally selected 20 images of animals to function as catch trials.

#### Similarity ratings and dimension extraction

We first obtained similarity spaces for the different object-knowledge types tested (visual information, functional information and manipulation information). We presented participants with words referring to our 80 manipulable objects and asked them to think about how similar these objects were in each of the knowledge types (e.g., function). We used an object sorting task to derive dissimilarities between our set of objects (74; Figure 1A), because sorting tasks have been shown to be a highly efficient and feasible way to obtain object similarities from large sets of objects. For each knowledge type, each participant was asked to sort all 80 objects into different piles such that objects in a pile were similar to each other, but different from objects in other piles, on the target knowledge type. Each participant went through the three knowledge types independently (order counterbalanced across participants) – that is, each participant was asked to judge the similarity of the objects 3 different times in a row (one for each knowledge type). We used words instead of pictures to avoid sorting based on exemplar-specific similarities. The sorting was performed in Microsoft power point. The final sorting was saved and later analyzed.

In Figure 1A, we show how data extracted from the piling task was used to produce dissimilarity matrices. Per participant, we obtained three (vision, function, and manipulation) 80 by 80 dichotomous (0 and 1) matrices that coded for membership of each pair of objects to the same sorting pile (i.e., if objects *i* and *j* were on the same pile, the value of the cell *ij* was 1, else it was 0). These individual matrices were then averaged over the participants and transformed into 3 final dissimilarity matrices – one per knowledge type.

The dissimilarity matrices were analyzed with the use of non-metric multidimensional scaling using Matlab. The number of dimensions to be extracted was determined based on *stress* value (Kruskal’s normalized stress 96) – i.e., the fit between the distances among the objects in the dimensional structure obtained with n-dimensions, and the scores in the input matrices. We obtained stress values for the MDS solutions with different numbers of dimensions. Stress values below 0.1 are considered acceptable (96, 136), suggesting that the dimensional solution, and estimated distances between objects, reasonably fits with the dissimilarities from the input matrix, while still imposing good dimensionality reduction. As such, we selected the first solution below stress values of 0.1 (see Figure S2 for scree plots for each of the knowledge types).

#### Label generation for extracted dimensions

To test for interpretability of the extracted dimensions, we asked participants to generate labels for the 15 key dimensions. Each participant went through all of the 15 key dimensions (order of dimensions randomized for each participant). For each dimension, participants were presented with the 20 most extreme objects of that dimension – 10 pertaining to each one of the extremes of the target dimension – rank ordered by the value on the target dimension. Pictures or words were presented such that in one side of the screen we lined up the 10 objects of one of the extremes, and on the other side of the screen the other 10 extreme objects of the dimension. At the center of the screen, we presented ellipsis between parenthesis to convey continuation and the presence of other objects in between. Participants were then told that they could generate up to 5 labels per dimension that, in their view, best explained the difference between the objects at the two extremes (see Figure 1B). Participants either saw images of the objects or words referring to the objects throughout the experiment. We used these two modalities of presentation to avoid presentation-specific results. We focused on which labels were more consensual across participants in the label generation task by collating the labels generated by the participants, and analyzing frequency of production of each label (see Figure S3 for the labels generated for each dimension).

#### Supervised learning of object-related dimensions

In this task, we taught participants to categorize a subset of our 80 objects in terms of their scores along a target dimension, and then tested whether their learning could be generalized to a subset of untrained (i.e., the remaining) items. We first divided the dimensions into 10 bins of 8 objects each, defined according to the values of the target dimension. We used the most extreme objects at each end of the dimension (i.e., those with the highest and the lowest score in the dimension) as anchors in the categorization tasks. Per trial, participants were asked to categorize, as fast as possible, the presented target objects (in a word format) as to whether they were closer to one or the other extreme object of the target dimension (e.g., pepper grinder or horn for the function dimension “Cooking vs. Sports”). To do so, participants had to press the right or left buttons of a button box – response assignment was randomized across participants. Participants were not told the labels of the target dimension, and were only informed as to whether the dimension was related to vision, function or manner of manipulation of the target objects.

The experiment was divided in three phases (in Figure 1C, we show the different phases of this experiment). In the first phase, we wanted participants to learn to associate objects with high or low dimension scores with the correct extreme. Thus, we selected 4 items from each of the 4 most extreme bins (i.e., bins 1, 2, 9, and 10), and 2 from each of the third most extreme bins (i.e., bins 3 and 8), in a total of 20 objects – i.e., we selected objects that were most strongly related with the two extremes of the dimension in order to establish a robust understanding of the target dimension. These were selected randomly within each bin per each participant. In each trial of the first phase of the experiment, participants were first presented with the words referring to the two extreme objects in the upper corners, according to the response assignment, and a fixation cross for 500 ms. We then presented the word referring to the target object at fixation, along with the words referring to the two extremes of the target dimension (in the upper corners of the screen). This remained on the screen for 2.5 s after which the correct response was presented right below the target object. Specifically, participants were presented with a sentence that said that the object presented was closer to one of the extremes. This sentence remained on screen for another 2.5 s. Participants were asked to pay attention to these sentences and learn the assignments between objects and extreme anchors. Each of the 20 objects was repeated three times in a total of 60 trials. Thus, in phase one, participants were not required to respond, but just to learn the associations of each of the presented objects with the extremes of the dimension.

In phase two, we wanted participants to continue learning the associations between target objects and the extreme anchors, and also to extended the learning set to all bins. Thus, we selected 5 items from each bin – including the 20 items used in phase one. In this phase, participants were presented again with the target object (3 repetitions, in a total of 150 trials), but this time were required to respond and categorize the target objects as being close to one of the extremes (e.g., closer to the pepper grinder or the horn). After responding, participants were given feedback as to whether they were correct or incorrect. The trial structure was in all similar to the one in phase 2 except that the object was presented for 2.5 s or until a response was obtained, and this was immediately followed by feedback as to whether the response given was correct. The feedback stayed on screen for 2 s.

Finally, in phase three, we wanted to test whether the learning could be generalized to untrained items. Thus, all items were used in this phase of the experiment (i.e., the 50 trained items and 30 untrained items – 3 from each of the ten bins) and were repeated 6 times (in a total of 480 trials). This phase of the experiment was in all equal to phase 2, except that there was not feedback given – that is, after categorizing a target object as to whether it was closer to one or the other extreme of the target dimension, participants would start the next trial.

All 15 key object-related dimensions were tested in this experiment (each participant as tested on only one dimension; 10 participants took part in the experiment for each dimension). Moreover, we added two control dimensions. In one of these controls, we took one of the real dimensions (the first function dimension) and randomly shuffled the scores of the dimension for the objects. For the other control, we took lexical frequency values (97) for each of the objects and rank ordered them in terms of these values. Each of these controls was run in 30 participants (i.e., 10 associated with each knowledge type). We used these dimensions to control for reliable generalization of object-related dimensional learning to untrained items.

To analyze the data of phase three, we first averaged responses per bin (3 untrained items repeated 6 times). For each participant and dimension, we plotted the percent responses towards the object with the highest score along that dimension as a function of the ten bins and fit a cumulative Gaussian to these data following the equation:

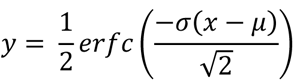

where y is the percent responses, x is percentile bins, σ is the slope of the cumulative Gaussian, μ is the midpoint of the curve, and erfc is the complementary error function. The cumulative Gaussian spans reports from 0% (lower asymptote) to 100% (upper asymptote). We expected that if the object-dimensions were cognitively important for our ability to process and recognize objects, then participants would be able to generalize their learning of the target dimensions to untrained items – thus, percent responses towards the extreme object with the highest score should increase as a function of the increase in the dimensional score per bin. R-square values per participant were obtained as a measure of goodness of fit between the cumulative Gaussian curve and each participant’s data. We used a liberal R-square cutoff value of 0.30. This specific cutoff is necessarily arbitrary, but we used this liberal criterion to allow us to determine if any given dimension is reasonably predictive of each participants’ behavior.

#### fMRI object categorization task

The fMRI task consisted of 1-3 sessions spread across separate days (due to the onset of COVID-19 pandemic, only 19 subjects completed all 3 sessions, with 5 & 2 remaining subjects completing 2 & 1 sessions, respectively). Each session contained 3 event-related runs (duration: 456 TRs (repetition time; 912 seconds), resulting in 3-9 completed runs per subject). In each run, subjects centrally fixated gray-scaled images of: 1) Manipulable objects (160 trials per run; 1 exemplar for each of the 80 objects was randomly selected from a larger stimulus set of 800 images (80 object identities x 10 exemplars each), and presented twice per run; and 2) ‘Catch’ animal stimuli (8 trials per run; 8 unique animal exemplars randomly drawn from a set of 20 images). Trial length was 4 seconds (2 seconds image presentation + 2 seconds fixation) and 54 null events (4 seconds fixation) were also included in the design. Because each object image was presented twice per run, randomization of trial order was performed for the first and second half of each run separately (i.e., to avoid strong recency effects where some items may, by chance, be repeated within a much smaller time period than others), where 80 object stimuli and 50% of catch trials and null events were presented in each half. Subjects were instructed to maintain fixation continuously (a fixation dot appeared in the center of the screen for the entirety of the run) and to make a simple button-press judgment for each image trial (object or animal).

#### MRI acquisition, preprocessing, and analysis

Scanning was performed with a Siemens MAGNETOM Prisma-fit 3T MRI Scanner (Siemens Healthineers) with a 64-channel head coil at the University of Coimbra (Portugal; BIN-National Brain Imaging Network). Functional images were acquired with the following parameters: T2* weighted (single-shot/GRAPPA) echo-planar imaging pulse sequence, repetition time (TR) = 2000 ms, echo time (TE) = 30 ms, flip angle = 75°, 37 interleaved axial slices, acquisition matrix = 70 x 70 with field of view of 210 mm, and voxel size of 3 mm^3^. Structural T1-weighted images were obtained using a magnetization prepared rapid gradient echo (MPRAGE) sequence with the following parameters: TR = 2530 ms, TE = 3.5 ms, total acquisition time = 136 s, FA = 7°, acquisition matrix = 256 x 256, with field of view of 256 mm, and voxel size of 1 mm^3^.

Data were preprocessed with SPM12 (i.e., slice-time correction, realignment (and reslicing), anatomical co-registration and segmentation, normalization, and smoothing). Analysis was performed on smoothed, normalized data (normalized to MNI template; 3mm isotropic voxels). General linear model (GLM) estimation was performed in SPM12 (data were high-pass filtered (256s) and a first-order auto-regression model (AR(1)) was used to estimate serial time-course correlations). For each subject, a GLM was estimated separately for each knowledge type because mathematically these dimensions are necessarily uncorrelated with each other (i.e., MDS produces uncorrelated dimensions). Each dimension/modulator was first scaled to an interval of 0-1 (due to the large differences in original scales of the key dimensions (ranging between-0.42 to 0.65) and then mean-centered, as is typical for parametric analysis (137). We ensured that serial orthogonalization of modulators was not implemented, so that all modulators would compete equally for model variance (rather than in the case of serial orthogonalization, assigning all shared variance to the first modulator, with subsequent modulators competing for the remaining unexplained variance, and therefore strongly biasing effects towards the first modulator, at the expense of all others; note that the lexical frequency dimension was also included as a modulator and as such, key dimensions only account for variance not explained by lexical frequency). The following regressors comprised the design matrix for a given run: 1 regressor for manipulable object stimuli (i.e. box-car regressor for all tool stimulus events, convolved with the SPM canonical haemodynamic response function), N key dimension modulators (e.g. vision dimensions 1-5), 1 lexical frequency modulator, 1 catch stimuli regressor (all catch stimuli; modeled but not analyzed), N ‘placeholder’ modulators for the catch regressor (i.e. a matched number of dimension modulators were required here for design balance, but these had no modulatory effect as all values were set to zero), 6 head-motion regressors (plus an intercept regressor at the end of the full design).

The resulting beta maps for each of the dimensions describe the extent to which object stimulus responses vary as a function of item position along a particular dimension. Importantly, because the obtained dimensions reflect an arbitrary directional ordering of items, and therefore are not inherently uni-directional (i.e., across repeated non-metric MDS solutions (with random initializations), relative item-to-item distances are stable but the overall item ordering can be reversed in either direction), both positive and negative slope effects here reflect sensitivity to a given dimension, based on either possible directional ordering. Thus, contrast images per dimension (contrast vector with 1 for the dimension modulator, 0 for all other regressors) were then entered into a group-level F-test to accommodate 2-tailed effects (i.e., positive or negative slopes that differed from zero). F-maps were FDR cluster-corrected (p<.05) with a height-threshold of p<.001 (but due to limited coverage for the first function dimension and fourth vision dimension, a slightly more lenient height-threshold of p<.005 (FDR cluster-corrected p<.05) was used, as previously motivated (e.g., 138–140). In short, the resulting F-maps show which brain areas demonstrate modulation sensitivity to each specific dimension.

For easy visualization of brain coverage associated with the dimensions of a particular knowledge type (see Figure 4), a composite dimension map was generated where supra-threshold voxels for each dimension were coded with a unique integer/color (and voxels with overlapping coverage from 2+ dimensions were coded with a different integer/colored black) Similar composite maps for the first 2 dimensions of each knowledge type were also created (see Figure 5). All maps were corrected for multiple comparisons (cluster-forming threshold p<.001 – or p<.005 for “Cooking vs. Sports” and “Material properties” – and FDR correction threshold p<.05). All maps were projected to an inflated surface with the CONN toolbox (141).

## Acknowledgements

This work was supported by the European Research Council (ERC) under the European Union’s Horizon 2020 research and innovation programme Starting Grant number 802553; “ContentMAP’’ to JA. AF is supported by a grant from the Biotechnology and Biology research council (BBSRC, grant number: BB/S006605/1) and the Bial Foundation, Bial Foundation Grants Programme Grant ID: A-29315, number: 203/2020, grant edition: G-15516. SK is supported by a Fundação para a Ciência e Tecnologia (FCT) Doctoral Grant SFRH/BD/145218/2019. DV is supported by a FCT Doctoral Grant SFRH/BD/137737/2018. FB is supported by a FCT Individual grant CEECIND/03661/2017. JW is supported by a FCT Individual grant CEECIND/03185/2021. The authors wish to thank Bradford Mahon for his comments on an earlier draft.

## SUPPLEMENTARY MATERIAL FOR

This section includes two supplementary tables and five supplementary figures:

**Table S1.**
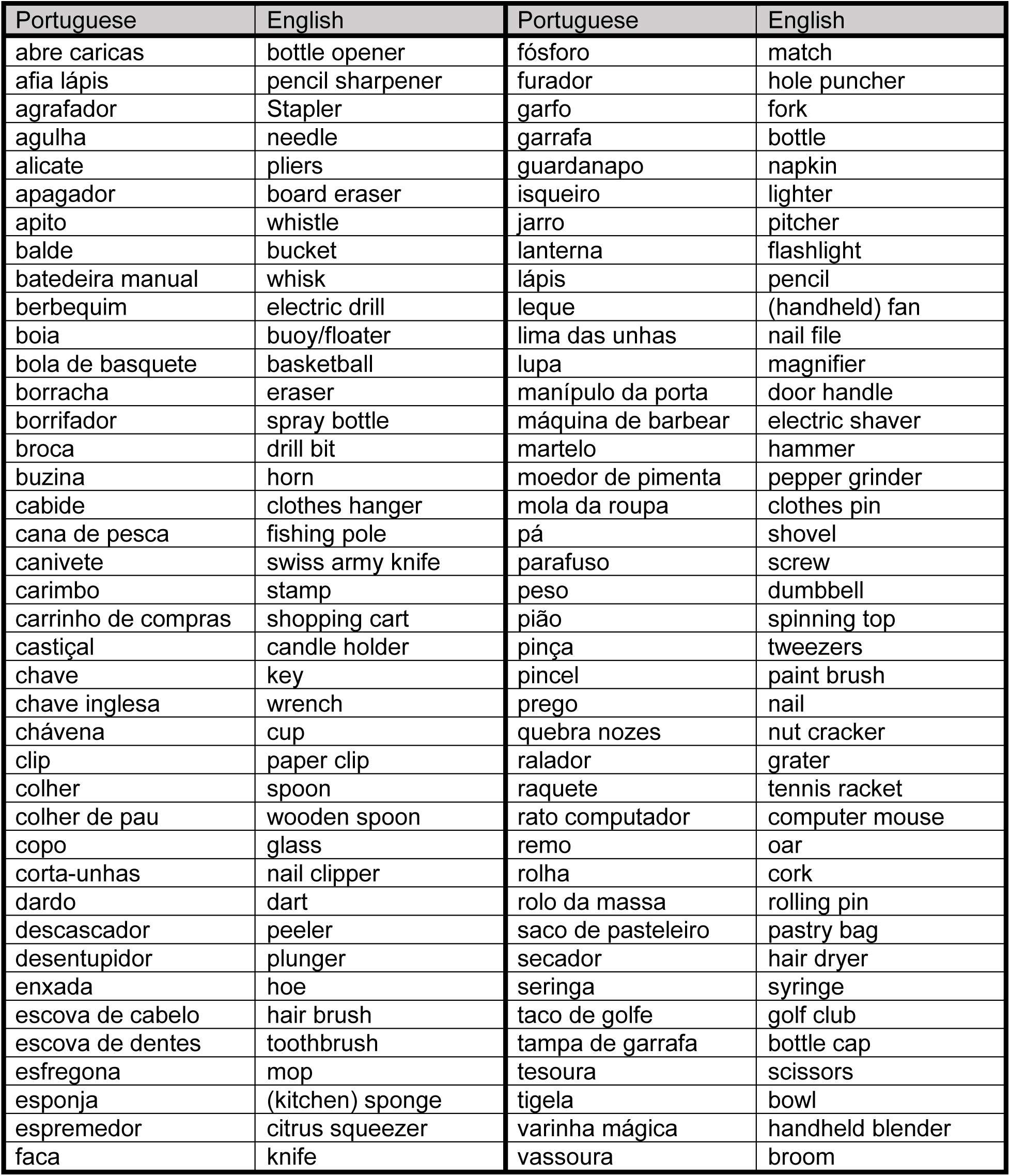
Items used in all of the experiments.

Here we present the original items used (in Portuguese) as well as the English translation of the Portuguese words used.

**Figure S1.**
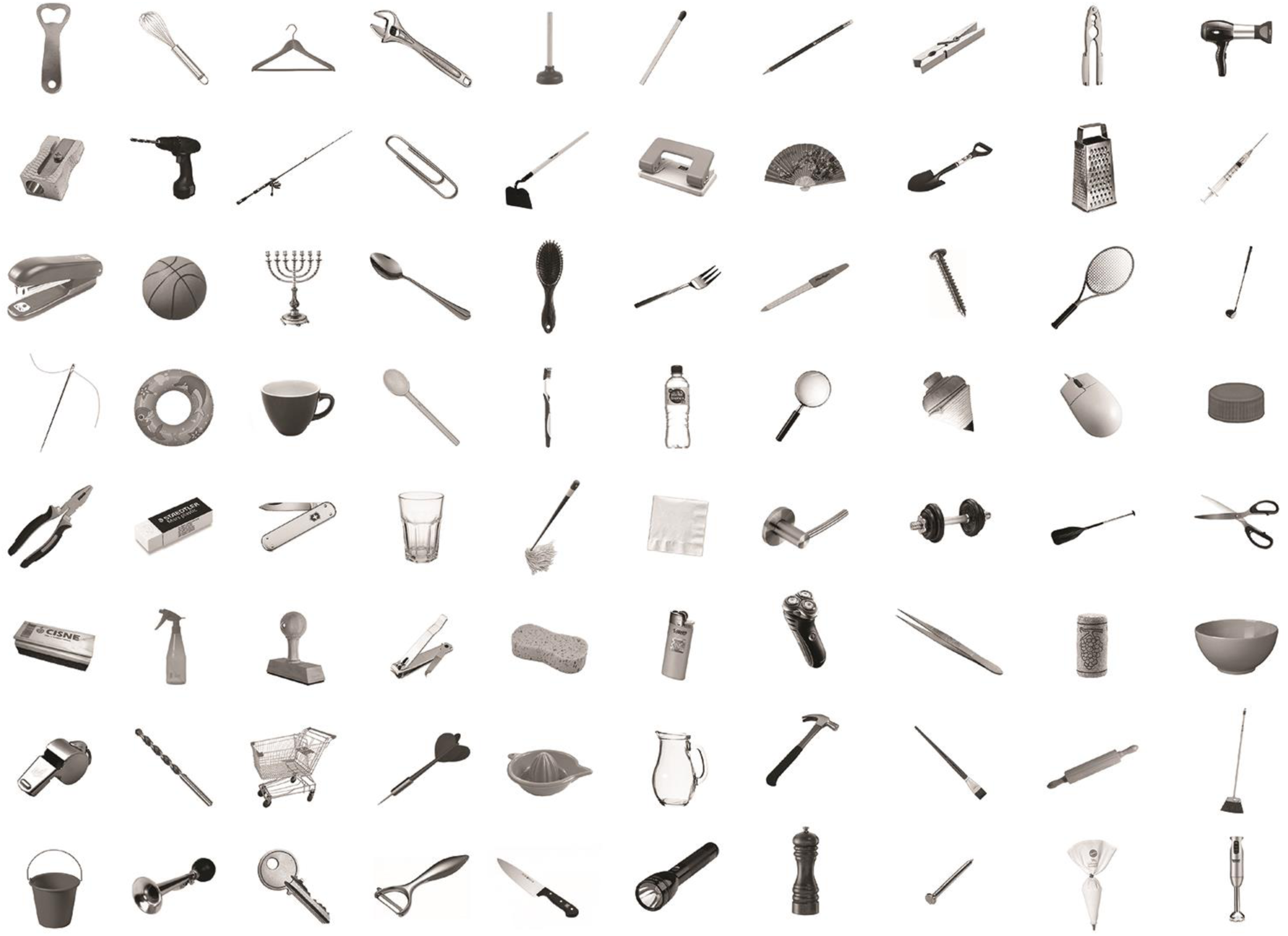
Examples of the 80 selected manipulable objects. Here, we present one of the ten exemplars for each of the 80 manipulable objects used in Experiment 4.

**Figure S2.**
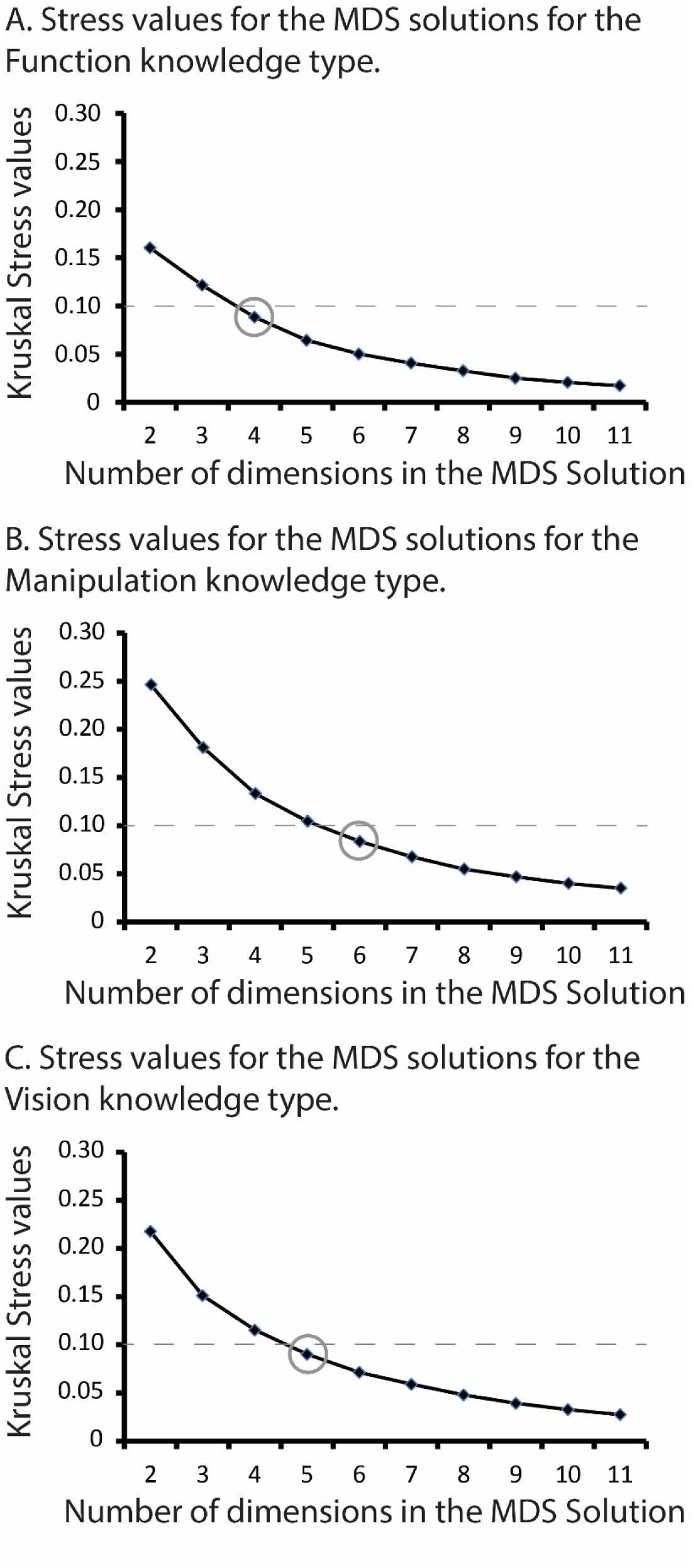
Stress values for the non-metric MDS analysis per each knowledge type. Here we show Kruskal Stress values for the different dimensional solutions for **A)** Function; **B)** Manipulation; and **C)** Vision. We chose the solutions with a stress value right below 0.1.

**Figure S3.**
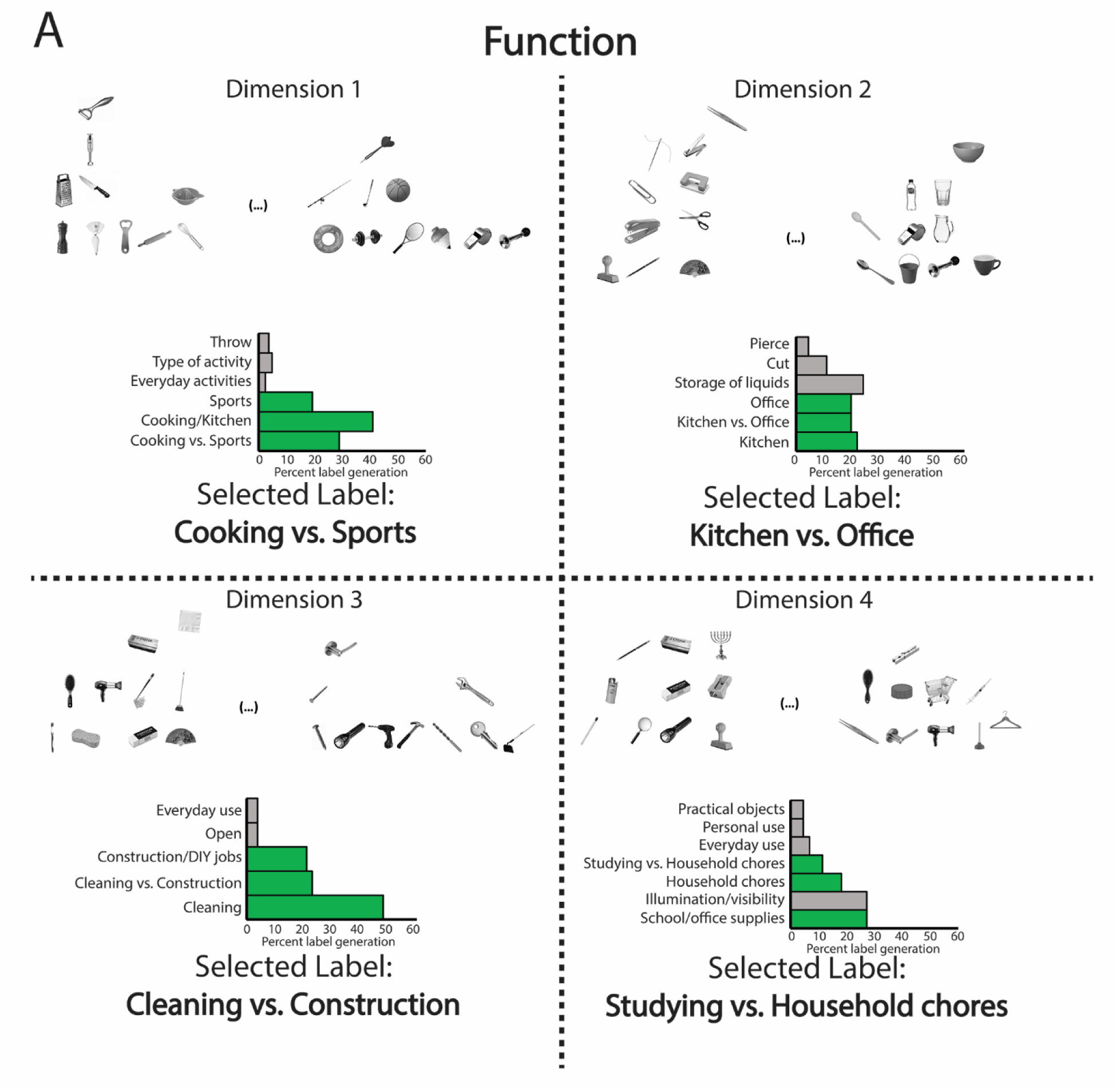
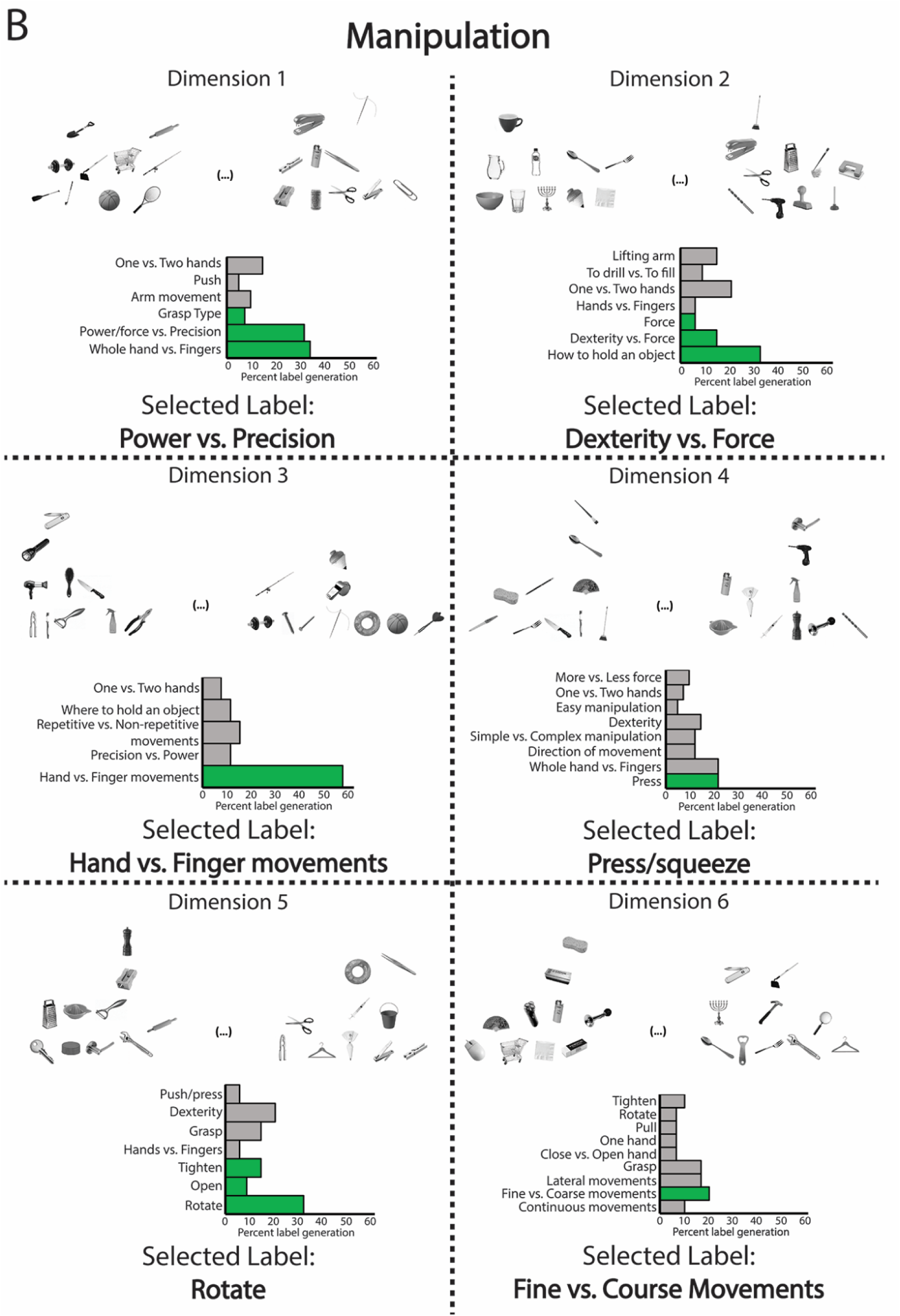
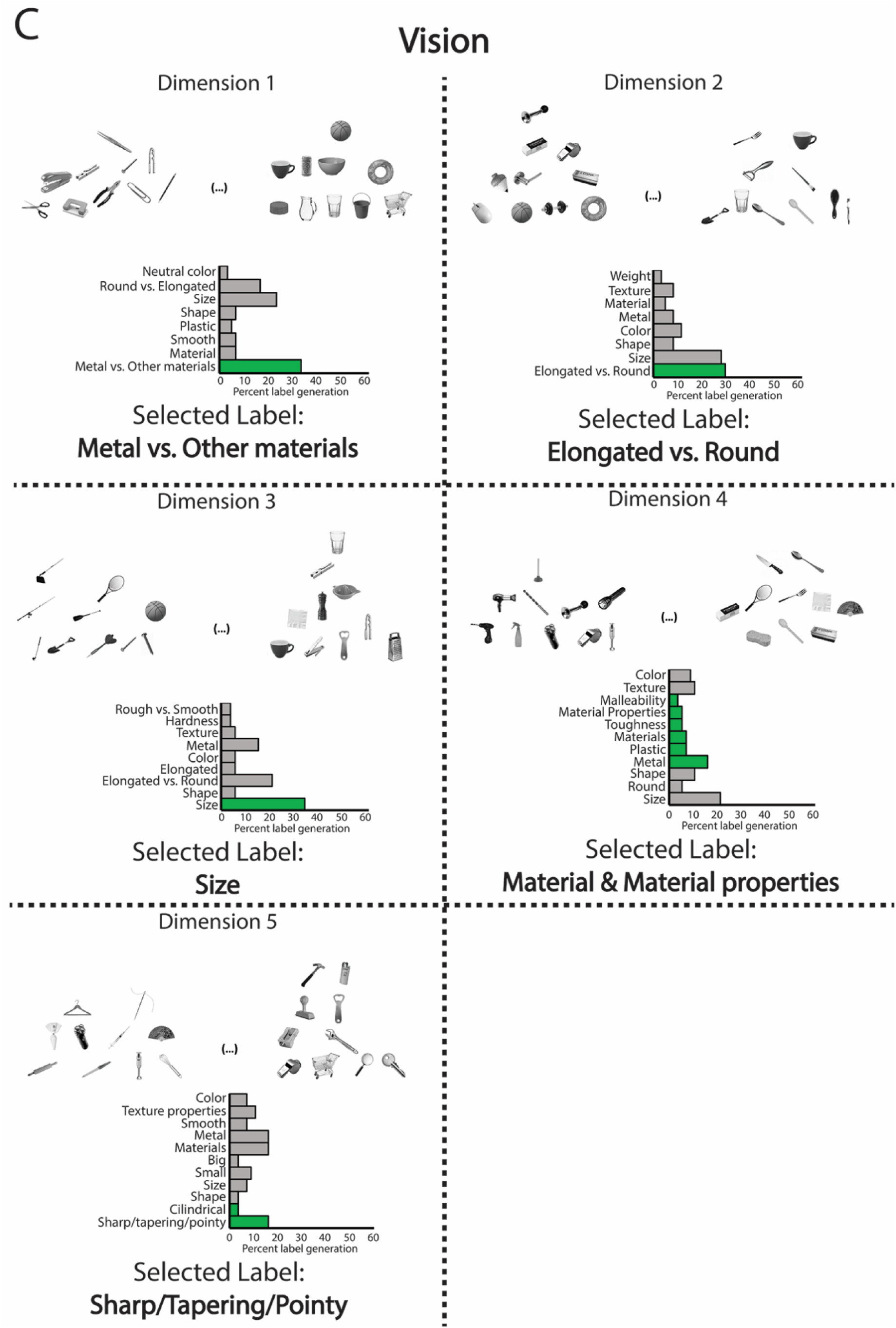
Labels generated per dimension. Here, we show percent label generation per dimension, along with an example of the images presented to the participant, for A) function; B) Manipulation; and C) Vision. Labels presented are collated from individual labels generated. Bars in green correspond to the labels that were put together as the winning label.

**Figure S4.**
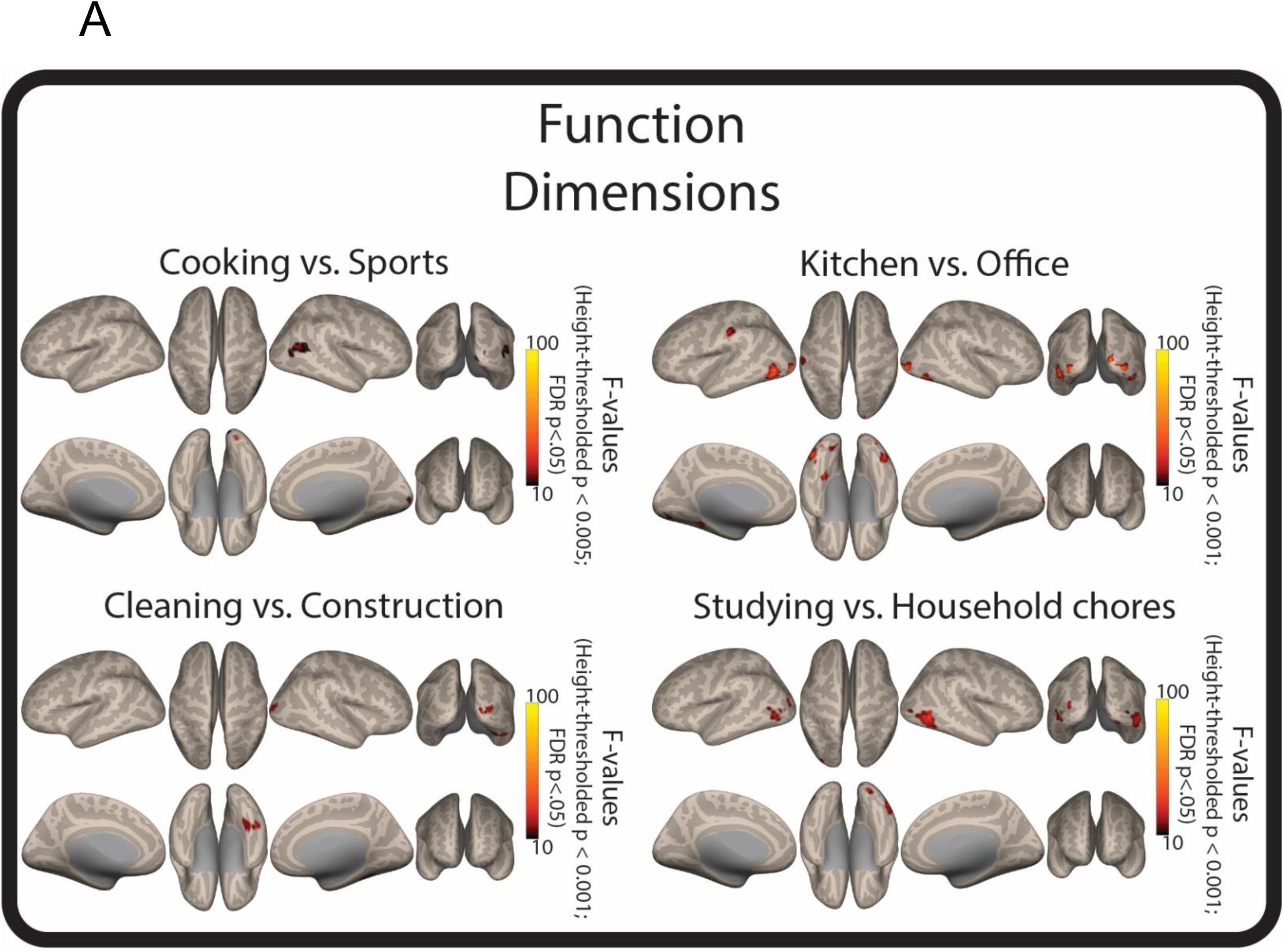
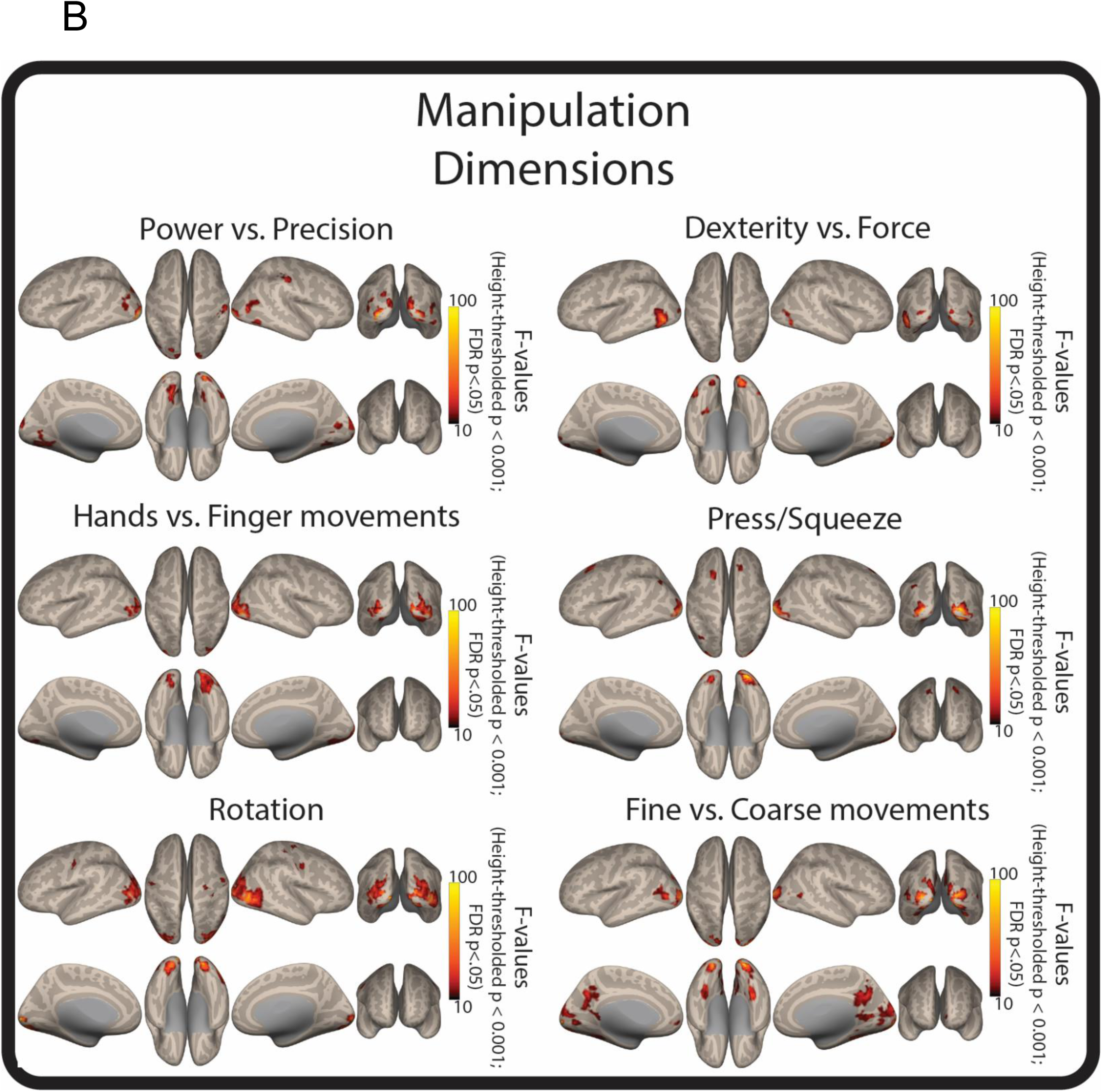
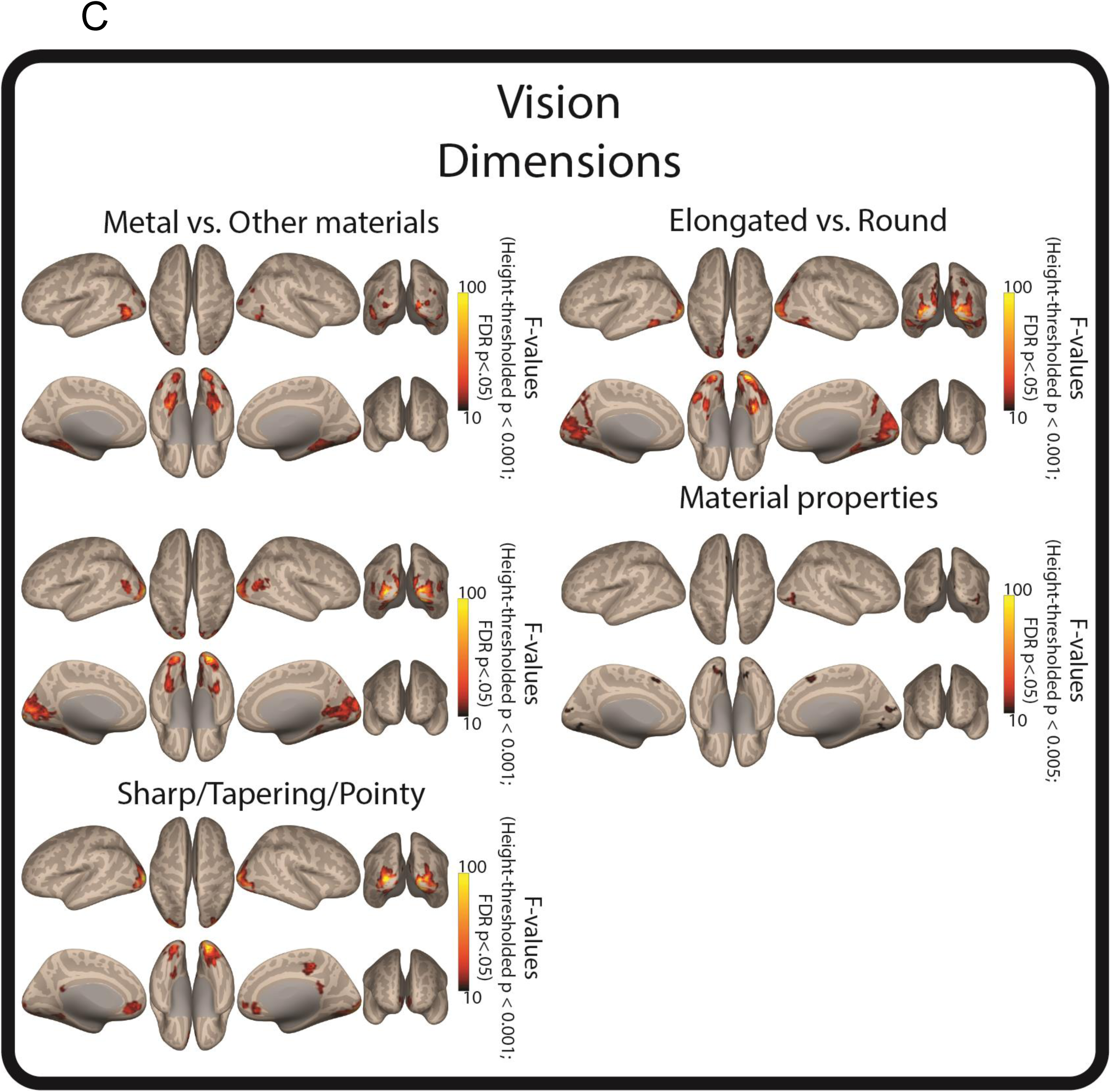
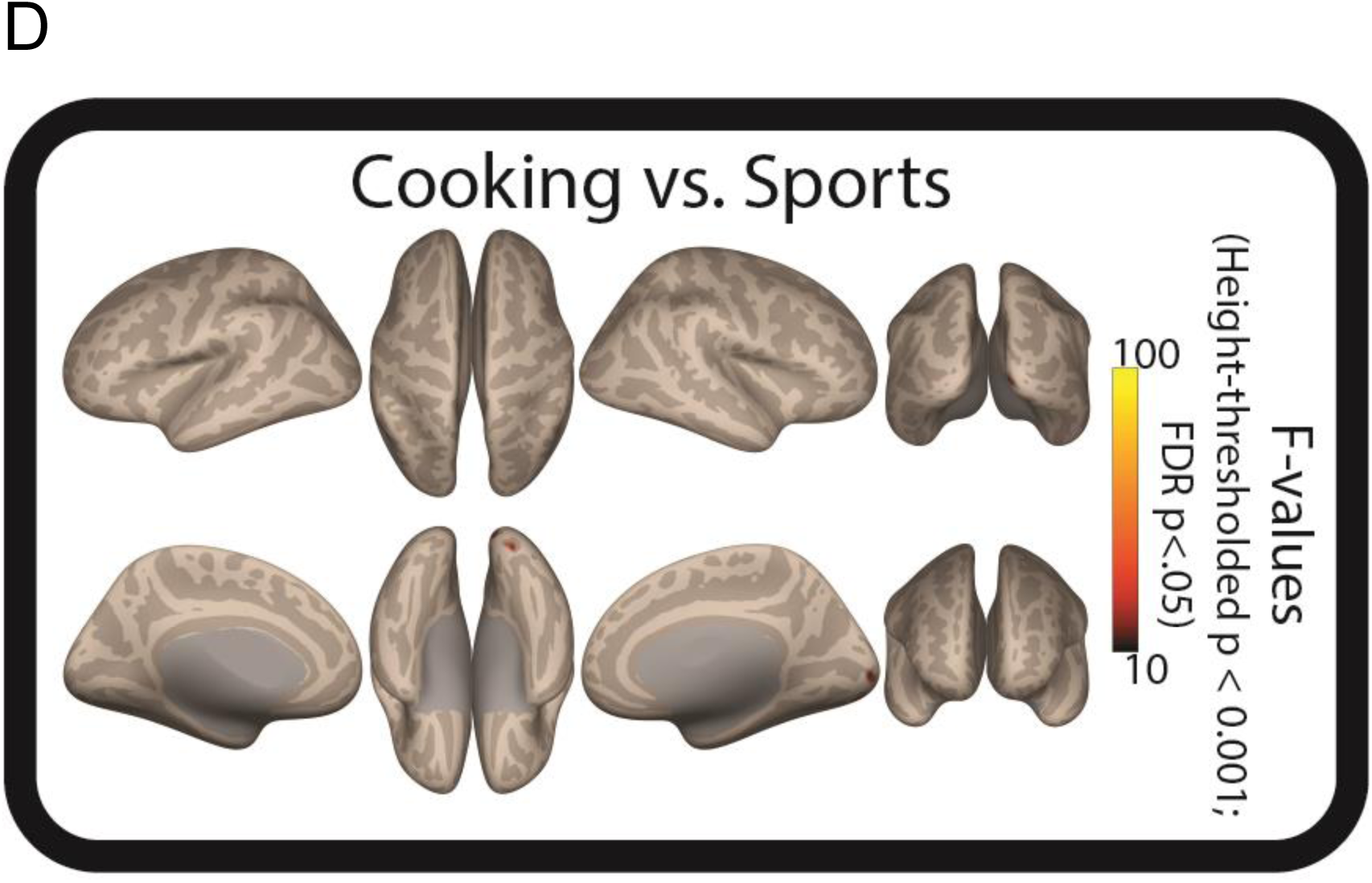
Neural effects of the object-related dimensions against 0 (i.e., against no modulation). Here we show dimension-specific F-maps against 0 for A) Function; B) Manipulation; and C) Vision. In D), we also show F-maps for “Cooking vs. Sports” cluster-forming height-thresholded at p<.001 and corrected at FDR p<.05. No cluster survived at this cluster-forming threshold for the dimension “Material properties”.

